# Brain-wide Cerebrospinal Fluid Recirculation Collapse in Alzheimer’s Disease

**DOI:** 10.64898/2026.10.01.755971

**Authors:** Moonseok Choi, Kyu-Sung Kim, Ii-woo Kim, Min Kyo Jung, Do-Geun Kim

## Abstract

Cerebrospinal fluid (CSF) flow and the glymphatic system are essential for maintaining brain homeostasis, yet the precise mechanisms driving their profound failure in Alzheimer’s disease (AD) remain largely elusive. Here, we reveal that macroscopic CSF propagates dorsally via the perivascular spaces (PAS) of major ventral arteries—the Circle of Willis (CW), middle cerebral artery (MCA), and olfactory artery (OlfA)—which undergo severe structural degradation in an AD mouse model. High-resolution imaging of the dorsal cortex demonstrates that this structural collapse induces a severe influx blockade; in AD, CSF tracers form massive aggregates within the PAS, physically decoupling the gliovascular interface, while tracers entering the parenchyma are pathologically sequestered by amyloid plaques. Furthermore, we identify an active, cholesterol-mediated clearance mechanism responding to exogenous CSF tracer infiltration. While healthy brains rapidly synthesize and efficiently clear this cholesterol via vascular efflux within 120 minutes, the AD model exhibits blunted cholesterol mobilization and a complete failure of capillary efflux. This terminal bottleneck forces uncleared cholesterol to retrogradely accumulate into pathological aggregates within perivascular pericytes and neuronal somata. Collectively, our findings demonstrate that AD dismantles the entire CSF recirculation loop—from macroscopic ventral entry to terminal microvascular efflux—highlighting pericytic and neuronal lipidosis as fundamental drivers of brain fluid stagnation.

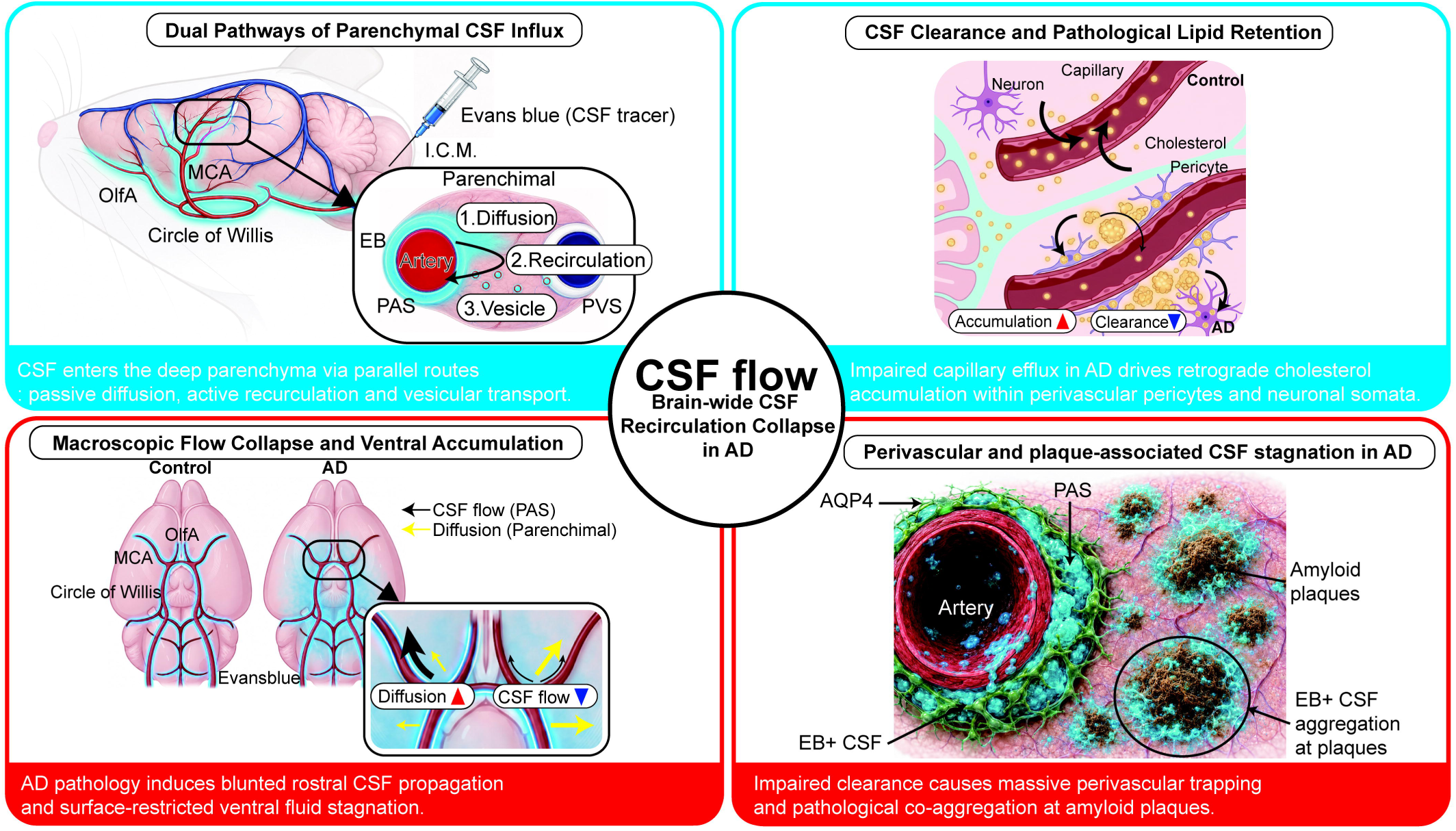

## Introduction

The exchange between cerebrospinal fluid (CSF) and interstitial fluid (ISF) is fundamental for supplying nutrients from the vasculature to diverse neural cells and for clearing metabolic waste generated by brain activity, thereby maintaining central nervous system homeostasis (1, 2). Beyond basic neuroprotection, the importance of this fluid dynamics—conceptually framed as the glymphatic system—has been increasingly recognized for its role in flushing out molecules ranging from small solutes to large protein aggregates through multiple paravascular, paravenous, and lymphatic routes (3, 4). To date, extensive research has demonstrated that periarterial CSF influx is tightly modulated by various physiological states; arterial pulsations driven by the cardiac cycle, sleep-wake transitions, and norepinephrine-mediated cortical arousal all significantly influence influx efficiency (5, 6). Furthermore, the structural configuration of the extracellular matrix (ECM), governed by the expansion and contraction of astrocytic networks, as well as the blood-brain barrier (BBB) fortified by astroglial aquaporin-4 (AQP4) water channels, act as critical gatekeepers for fluid entry (6, 7). However, despite these insights, the comprehensive spatiotemporal mapping of brain-wide CSF flow—specifically, how fluid propagates along major basal vessels from the ventral to dorsal regions over time, and the exact physical nature by which it infiltrates the deep parenchyma—remains poorly understood.

Crucially, recent mathematical advection-diffusion models suggest a fundamental limitation in the current paradigm: passive diffusion alone cannot fully account for the rapid and deep parenchymal transport of CSF from the perivascular space (PAS) to the ISF, and ultimately to the draining veins(8, 9). These models indicate that parenchymal transit likely requires an active, yet-to-be-identified structural or packaging system to facilitate macromolecular delivery, a hypothesis that traditional diffusion-centric models fail to address. Furthermore, while previous imaging studies have largely focused on dorsal or lateral cortical surfaces, the macroscopic fluid dynamics within the ventral base of the brain—where major cerebral arteries such as the Circle of Willis (CW), middle cerebral artery (MCA), and olfactory artery (OlfA) branch and distribute—have been remarkably underexplored (10, 11). Consequently, the ultimate fate of the CSF-ISF mixture after it enters the parenchyma, how it interacts with neuronal somata, and the precise trajectories by which it eventually re-enters the vascular compartment to complete a recirculation loop, remain critical knowledge gaps.

Understanding this full circulation loop is essential, particularly because the progressive failure of CSF transport and glymphatic clearance is increasingly implicated as a core pathology in neurodegenerative disorders such as Alzheimer’s disease (AD) (12). Pathological studies reveal that exposure to amyloid-beta (Aβ) induces severe neurovascular dysfunction, impairs macrophage function, and provokes the loss of perivascular AQP4 polarization, thereby suppressing periarterial CSF influx and ISF clearance (13, 14). In both clinical settings and AD mouse models, advanced imaging modalities, such as dynamic contrast-enhanced magnetic resonance imaging (DCE-MRI), consistently report profound reductions and spatial rerouting in global CSF flow (15, 16). Yet, these observations merely report the consequence—a global flow reduction—without pinpointing the exact anatomical and microstructural bottlenecks. Specifically, it remains unknown how AD pathology alters ventral dynamics at the major arterial roots (Circle of Willis, MCA, OlfA), how deep parenchymal fluid exchange and vascular re-entry fail, and what precise physical role Aβ plaques play in disrupting this delicate fluid architecture.

To resolve these enduring black boxes in brain fluid dynamics and to elucidate the structural drivers of clearance failure in AD, we employed a multiscale imaging platform. Utilizing the 5xFAD mouse model of amyloidosis, we tracked the spatiotemporal progression of intra-cisterna magna (I.C.M.) injected Evans Blue (EB) tracer. By integrating macroscopic whole-brain optical mapping of cleared tissues with high-resolution confocal section imaging and super-resolution tissue Correlative Light and Electron Microscopy (CLEM), we map the trajectory of CSF from its initial ventral periarterial distribution into the deep parenchyma, and its eventual vascular return. Through this comprehensive approach, we characterize a brain-wide CSF recirculation loop, and dissect the exact macroscopic and microscopic pathological bottlenecks—including perivascular decoupling, parenchymal stagnation, and plaque-associated fluid trapping—that collectively dismantle brain fluid circulation in AD.

## Results

### Periarterial influx drives brain-wide CSF recirculation

While extensive research has investigated global CSF flow, the precise mechanisms by which CSF enters the PAS and navigates the dense brain parenchyma—interacting with neurons and the ECM— remain poorly defined (17). To delineate these dynamics, we employed a comprehensive multi-scale mapping strategy. Following I.C.M. administration of the EB tracer, we tracked CSF flow trajectories from macroscopic brain-wide distributions down to cellular and vesicular architectures using high-resolution confocal microscopy (Figure 1A).

**Figure 1.**
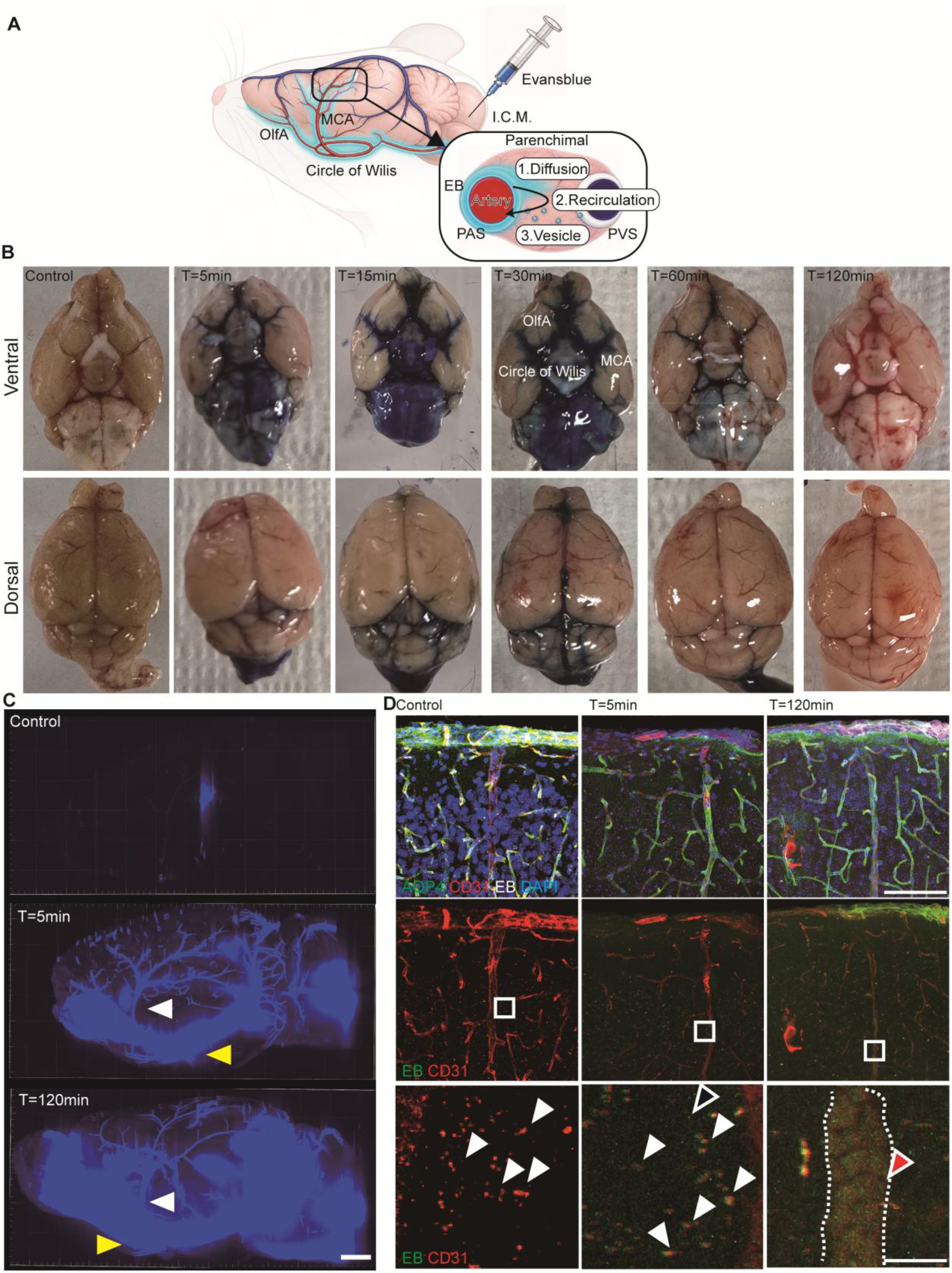
Multiscale imaging reveals brain-wide cerebrospinal fluid propagation, organized parenchymal transport, and delayed vascular return. **(A)** Schematic illustration of the experimental workflow mapping the brain-wide cerebrospinal fluid (CSF) recirculation pathway. Following intra-cisterna magna (I.C.M.) injection, the Evans Blue (EB) tracer rapidly distributes along the periarterial spaces (PAS) of major basal vessels, including the Circle of Willis, middle cerebral artery (MCA), and olfactory artery (OlfA). Within the parenchyma, CSF solutes transit toward the perivenous space (PVS) via a combination of passive diffusion and organized, vesicle-mediated transport architectures. **(B)** Representative optical images of the ventral and dorsal brain surfaces captured at baseline (Control) and at 5, 15, 30, 60, and 120 min post-I.C.M. EB injection. **(C)** Three-dimensional light-sheet fluorescence microscopy of tissue-cleared hemibrains, visualizing the global volumetric distribution of the EB tracer (blue) at baseline, 5 min, and 120 min post-injection. Yellow arrowheads highlight the robust accumulation of the EB tracer along the main arterial architectures at the ventral base. White arrowheads delineate the distinct propagation of the CSF tracer from the ventral origin ascending toward the dorsal surface along the MCA. **(D)** Representative high-resolution confocal images of the cerebral cortex at baseline, 5 min, and 120 min post-injection. Brain sections were immunolabeled for aquaporin-4 (AQP4, green) and CD31 (red), and counterstained with DAPI (blue); EB tracer fluorescence is represented in white (top row: scale bar 100um) and green (middle and bottom rows: scale bar 10um). At 5 and 120 min post-injection, diffuse EB signals are detected within the extracellular matrix (open arrowheads), alongside distinct CD31^+^ vesicular structures co-localizing with EB (white arrowheads). Crucially, at 120 min post-injection, EB tracer is observed re-entering the vascular compartment, localized within the lumen (red arrowheads) of CD31^+^ vessels (demarcated by white dashed lines). These combined observations indicate that organized, structure-mediated parenchymal transit ultimately leads to delayed vascular return, completing the brain-wide CSF recirculation loop.

Macroscopic optical imaging revealed that the EB tracer does not merely diffuse passively across the brain surface. Instead, it propagates rapidly along the PAS of major ventral arteries, predominantly the CW, MCA, and OlfA. This ventrally originated flow subsequently ascends toward the dorsal regions. Notably, under low-dose injection conditions, the tracer is almost entirely cleared systemically by 120 minutes post-injection, highlighting the highly dynamic nature of this clearance system (Figure 1B). To overcome the resolution limits of surface optical imaging and fully capture global volumetric dynamics, we performed three-dimensional light-sheet fluorescence microscopy on tissue-cleared hemibrains. These 3D reconstructions confirmed that even at 5 minutes post-injection, CSF flow is not static but highly active, with the EB signal robustly ascending along the MCA to reach dorsal territories. By 120 minutes, this tracer movement continues along the vasculature, achieving extensive penetration into the deep dorsal parenchyma (Figure 1C).

To resolve the micro-anatomical details of this parenchymal influx, we evaluated the dorsal cortex (layers 1–3) using high-resolution confocal microscopy of brain sections immunolabeled for vascular endothelium (CD31) and astrocyte endfeet (AQP4). At both 5 and 120 minutes post-injection, we observed two concurrent modes of parenchymal entry. Alongside the expected diffuse EB signal permeating the ECM (Figure. 1d, open arrowheads), we identified highly organized CD31^+^ vesicular structures co-localizing with EB (Figure 1D, white arrowheads). Importantly, these discrete CD31^+^ vesicular pathways distributed across the parenchyma and within neuronal somata were also observed in the baseline control group, indicating an endogenous structural network. Strikingly, at the 120-minute time point, the EB tracer that had entered the parenchyma via these diffusion and vesicular pathways was observed re-entering the lumen of major CD31^+^ arteries and capillaries—providing direct visual evidence of vascular recirculation (Figure 1D, red arrowheads).

Collectively, these multiscale observations indicate that the major ventral arteries and their associated PAS play a critical role in the initial brain-wide distribution of CSF. Any structural or functional impairment in this ventral network would likely severely suppress the subsequent dorsal propagation of CSF flow. Furthermore, once CSF (or ISF) infiltrates the parenchyma via the PAS, it undergoes tissue exchange through both passive diffusion and organized vesicular pathways. Crucially, in addition to the canonical efflux routes toward the venous system, these data reveal a previously unrecognized recirculation pathway wherein parenchymal fluid re-enters the arterial vasculature, completing a comprehensive brain-wide fluid loop.

### Structural periarterial bottlenecks and CSF flow collapse in AD

Numerous clinical and preclinical studies have established that AD is characterized by a global reduction in CSF flow, a deficit that impairs the clearance of Aβ and accelerates disease progression (12, 14, 15). Building upon our experimental scheme that the ventral CW, MCA, and OlfA act as the primary gateways for brain-wide CSF distribution, we investigated whether AD pathology directly disrupts these specific influx corridors and their subsequent dorsal propagation. Furthermore, we hypothesized that such structural stagnation at the ventral base would forcefully reroute fluid dynamics, promoting localized basal diffusion (Figure 2A).

**Figure 2.**
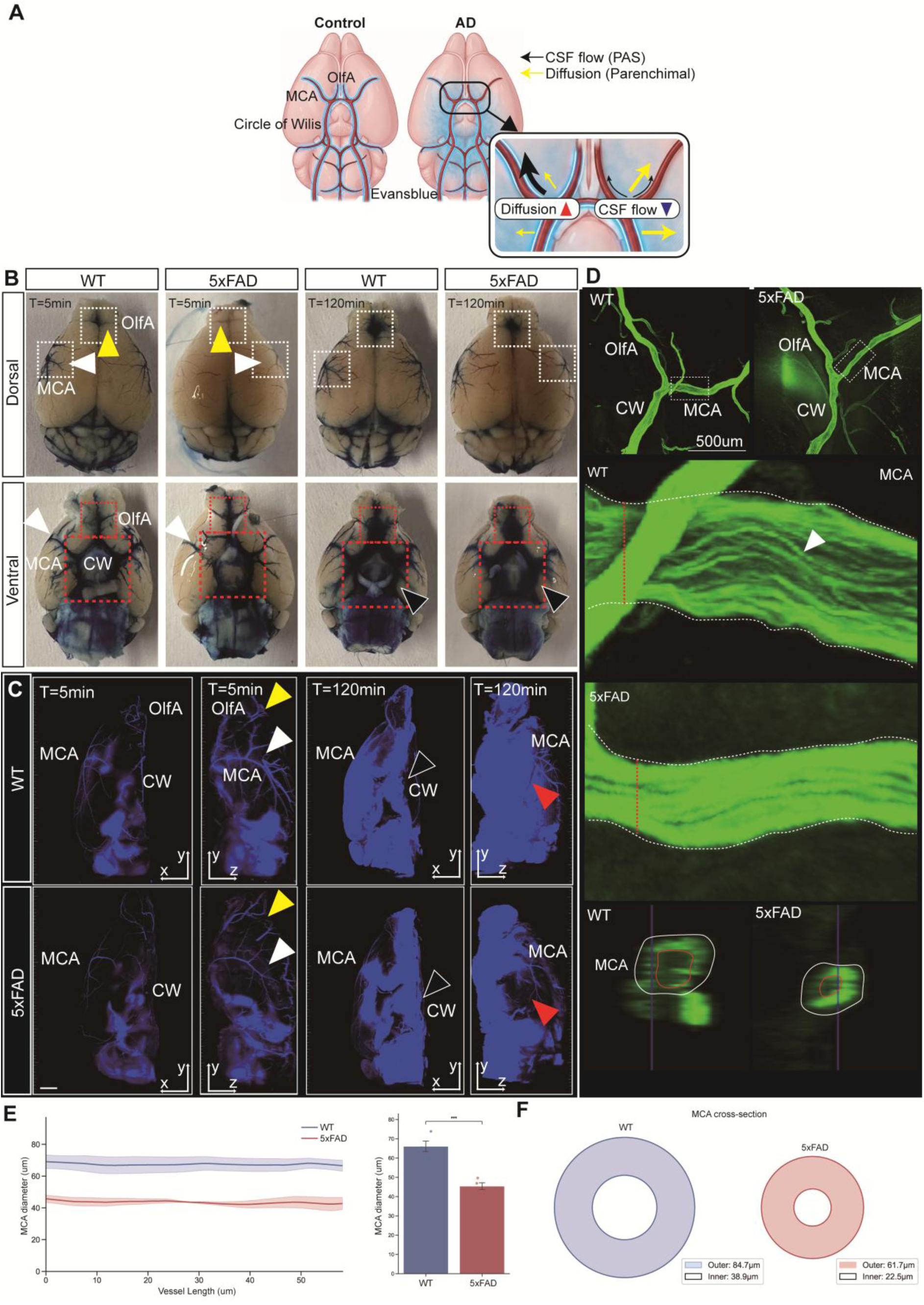
Alzheimer’s disease pathology drives macroscopic CSF flow collapse and structural periarterial bottlenecks. **(A)** Schematic illustration comparing normal cerebrospinal fluid (CSF) flow with the pathological flow collapse observed in Alzheimer’s disease (AD). Black arrows indicate the influx of CSF along the periarterial space (PAS), while yellow arrows represent the subsequent fluid diffusion into the deep parenchyma. In AD, profound basal stagnation and impaired parenchymal diffusion severely restrict brain-wide circulation. **(B)** Representative macroscopic optical images of wild-type (WT) and 5xFAD mouse brains at 5 min and 120 min post-intra-cisterna magna (I.C.M.) injection of Evans Blue (EB) tracer. Top (Dorsal view): White dashed boxes highlight the propagation of CSF along the middle cerebral artery (MCA, white arrowheads) and olfactory artery (OlfA, yellow arrowheads). Bottom (Ventral view): Red dashed boxes highlight the initial ventral influx networks, including the MCA (white arrowheads), OlfA, and Circle of Willis (CW, black arrowheads). The 5xFAD mice exhibit severe ventral pooling and an almost complete failure of dorsal CSF distribution. **(C)** Three-dimensional light-sheet fluorescence microscopy of tissue-cleared hemibrains at 5 min and 120 min post-injection (Scale bar, 1 mm). White, yellow, and black arrowheads denote the tracer signals along the MCA, OlfA, and CW, respectively. Red arrowheads highlight the deep parenchymal EB tracer penetration in the dorsal region, which is markedly diminished in the 5xFAD group. **(D)** High-resolution 3D reconstruction (top, middle) and cross-sectional imaging (bottom) of the ventral CW, OlfA, and MCA regions in cleared hemibrains (Scale bar, 500 um). Middle: Zoomed-in views of the MCA. White dashed lines delineate the outer boundary of the PAS. Strikingly, the EB signal does not flow as a uniform cylindrical sheet within the PAS; rather, it propagates along multiple distinct, parallel linear tracks (white arrowheads). This highly organized linear flow pattern remains present in 5xFAD mice, but the overall MCA diameter is visibly restricted. Bottom: Cross-sectional profiles corresponding to the red dashed lines. The inner red solid lines indicate the vascular lumen (displaying weak EB signal), while the outer white solid lines demarcate the EB-rich PAS boundary. **(E)** Quantitative spatial profiling of the MCA PAS boundary. Left: The continuous distance profile of the MCA outer diameter demonstrates a profound structural narrowing in 5xFAD mice compared to WT. Right: Bar graph representing the mean MCA overall diameter, revealing a statistically significant constriction in the 5xFAD group. **(F)** Schematic geometric representation of the MCA cross-sectional dimensions derived from the imaging data, illustrating that both the inner vascular lumen diameter and the outer PAS boundary are substantially reduced in the 5xFAD model, establishing a critical anatomical bottleneck for CSF influx. Data are presented as mean ± S.E.M. (*P < 0.05, P < 0.01).

To test this, we compared the macroscopic spatiotemporal propagation of I.C.M.-injected EB tracer in wild-type (WT) and 5xFAD mice. In WT controls, dorsal optical imaging at 5 minutes post-injection revealed clear, directed tracer propagation along the MCA (Figure 2B, Dorsal, white arrowheads) and OlfA (Figure 2B, Dorsal, yellow arrowheads). In stark contrast, 5xFAD mice exhibited a drastic reduction in tracer arrival at these dorsal territories. By 120 minutes, while the WT brain demonstrated robust and continuous dorsal tracer accumulation, the 5xFAD group showed an almost complete absence of dorsal EB signal (Figure 2B, Dorsal). Intriguingly, observing the ventral surface revealed a distinct pathological flow pattern. In WT mice, initial tracer influx branched symmetrically from the CW into the bilateral MCAs. However, 5xFAD mice frequently displayed a dramatic left-right flow asymmetry, with tracer propagation severely blunted on one side (Figure 2B, Ventral, white arrowheads). By 120 minutes, WT mice maintained directed tracer movement along the vascular tracts accompanied by localized tissue diffusion. Conversely, in the 5xFAD group, directed upward vascular propagation was markedly diminished; instead, the tracer appeared to stall and diffuse abnormally outward around the ventral CW, MCA, and OlfA, creating a hazy, superficial accumulation indicative of severe basal fluid pooling (Figure 2B, Ventral, red dashed boxes and black arrowheads).

To volumetrically confirm this spatial redistribution, we utilized three-dimensional light-sheet microscopy of tissue-cleared hemibrains (Figure 2C). Consistent with the surface imaging, lateral projections at 5 minutes revealed a profound reduction in CSF ascending the MCA (Figure 2C, white arrowheads) and OlfA (Figure 2C, yellow arrowheads) in 5xFAD mice compared to WT. By 120 minutes, the tracer in WT brains exhibited sharp, well-defined diffusion patterns ascending from the CW. In the 5xFAD model, however, the EB signal around the CW was abnormally broadened, further corroborating the basal fluid stagnation (Figure 2C, 120 min left, black arrowheads). Moreover, 120-minute lateral views definitively demonstrated that while the WT tracer successfully permeated the deep dorsal cortex, the 5xFAD tracer remained densely trapped at the ventral base, completely failing to reach the dorsal parenchymal territories (Figure 2C, 120 min right, red arrowheads).

We hypothesized that this profound failure of directed CSF flow stems from structural abnormalities within the major ventral arterial networks themselves. To resolve the fine architecture of these tracts, we employed high-resolution intravital microscopy (IVIM) to visualize the ventral CW, MCA, and OlfA (Figure 2D, top). Detailed magnification of the MCA and its surrounding PAS revealed a striking, previously unrecognized flow architecture: the EB tracer did not flow as a uniform, homogeneous cylinder filling the PAS, but rather propagated along multiple distinct, parallel linear tracks within the perivascular boundary (Figure 2D, middle, white arrowheads). While this linear multi-track organization was preserved in 5xFAD mice, the overall dimensions of the MCA and its PAS were significantly constrained.

Quantitative spatial profiling along the vessel length confirmed that the mean MCA diameter— measured continuously as the EB-labeled channel width in longitudinal IVIM sections—was significantly reduced in the 5xFAD group (46.65 μm) compared to WT (66.09 μm) (Figure 2E). Independently, cross-sectional analyses at defined points along the vessel demonstrated that the EB signal was strictly confined to the outer PAS domain rather than the vascular lumen, and that the pathological constriction in 5xFAD mice affected both the inner vascular lumen (WT: 38.9 μm vs. 5xFAD: 22.5 μm; red solid lines) and the outer PAS boundary (WT: 84.7 μm vs. 5xFAD: 61.7 μm; white solid lines) (Figure 2D bottom, Figure 2F). Because the PAS constitutes an annular channel bounded internally by the vascular wall and externally by the AQP4 endfoot layer, we defined the effective PAS width as the difference between these outer and inner boundaries (WT: ∼45.8 μm; 5xFAD: ∼39.2 μm), which we use below to approximate the hydrodynamic consequence of this structural narrowing.

Collectively, these findings indicate that the macroscopic collapse of brain-wide CSF circulation in AD is heavily driven by critical structural bottlenecks at the ventral base of the brain. The physical narrowing of the CW, MCA, and OlfA arterial and perivascular corridors effectively chokes off fluid influx, shifting the overall brain fluid dynamics from directed dorsal propagation to pathological ventral stagnation and basal diffusion. This macroscopic constriction of the ventral periarterial corridors strongly supports recent evidence that morphological alterations within these periarteriolar spaces are a primary anatomical determinant of altered glymphatic flow dynamics and brain-wide CSF transport deficits in amyloid pathology (18, 19).

### Parenchymal bottlenecks and CSF recirculation failure

Having established that structural bottlenecks at the ventral base severely restrict macroscopic fluid distribution, we next examined how AD pathology alters microscopic fluid dynamics within the dorsal cortex. As a conceptual framework, we outline this microscopic collapse as a cascading failure: structured parenchymal entry is impaired, leading to massive perivascular stagnation, subsequent fluid entrapment by amyloid plaques, and the ultimate disruption of the vascular recirculation loop (Figure 3A). High-resolution confocal imaging of cortical layers 1–3 (labeled for CD31, AQP4, and EB) revealed a profound disruption at the gliovascular interface. In WT mice, the CSF tracer moved seamlessly through the PAS and into the parenchyma without aberrant accumulation at both 5 and 120 minutes post-injection (Figure 3B). Conversely, 5xFAD mice exhibited massive EB aggregations (∼50 µm in diameter) trapped between the CD31^+^ endothelial layer and the AQP4^+^ astrocytic endfeet as early as 5 minutes, a severe perivascular stagnation that persisted through the 120-minute time point (Figure 3B and 3C).

**Figure 3.**
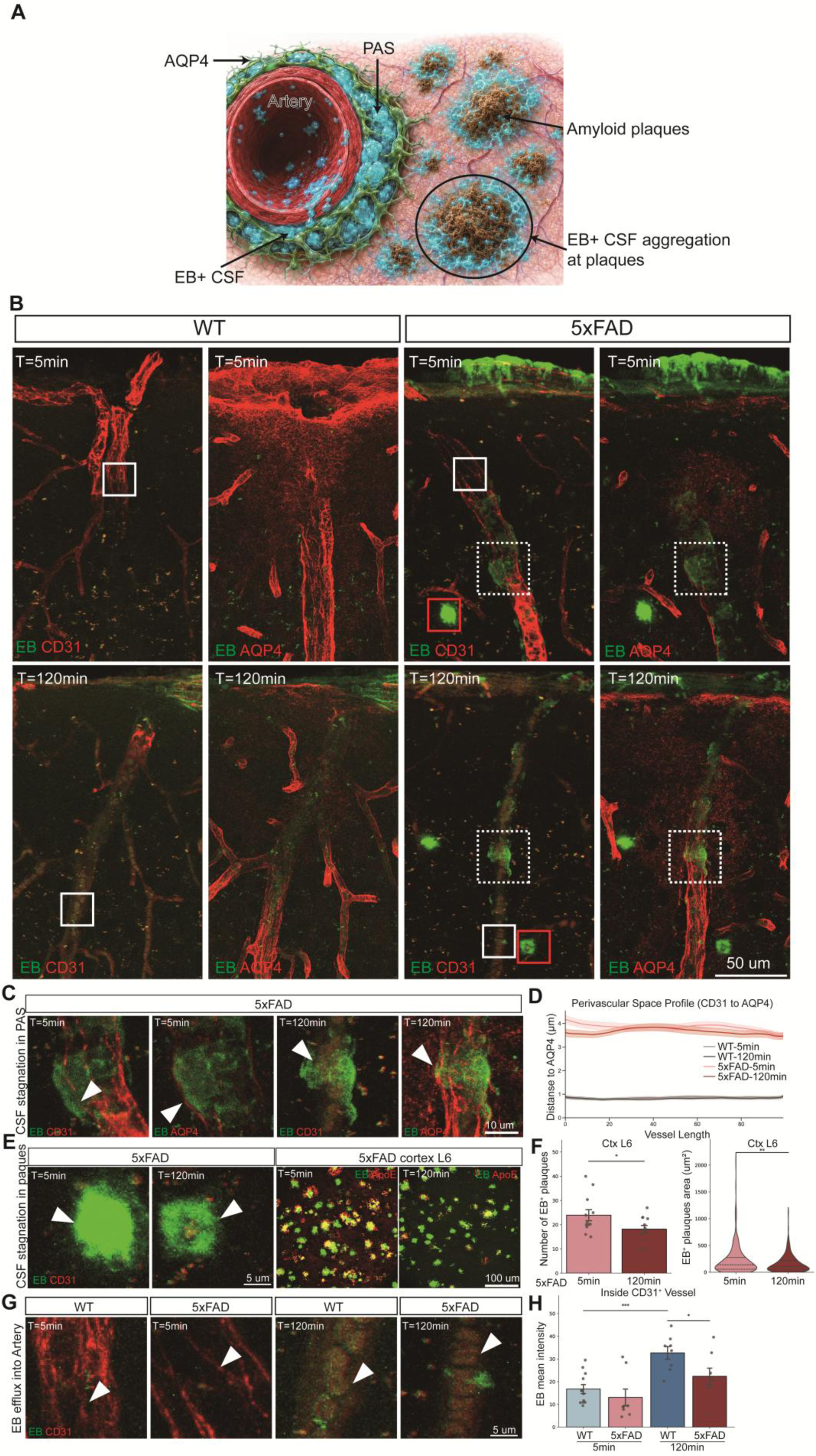
Alzheimer’s disease pathology induces perivascular stagnation, gliovascular decoupling, and the failure of vascular recirculation. **(A)** Schematic illustration summarizing the cascade of microscopic fluid dynamics collapse in the Alzheimer’s disease (AD) brain. In the pathological state, structured cerebrospinal fluid (CSF) influx from the periarterial space (PAS) into the parenchyma is severely impaired. Consequently, CSF (cyan) undergoes massive stagnation within the PAS, physically restricted by the astrocytic aquaporin-4 (AQP4) boundary. Furthermore, fluid that manages to diffuse into the tissue becomes abnormally entrapped around amyloid plaques, ultimately disrupting the completion of the vascular recirculation loop. (B) Representative confocal micrographs of major arteries in the dorsal cortex (layers L1 and L2/3) of wild-type (WT) and 5xFAD mice at 5 min and 120 min post-intra-cisterna magna (I.C.M.) injection of Evans Blue (EB) tracer. (C) Magnified views (from dashed white boxes in b) highlighting acute CSF stagnation within the PAS in 5xFAD mice. Unlike WT, 5xFAD mice exhibit robust, abnormal EB accumulation (white arrowheads) bounded by the AQP4+ astrocytic endfeet as early as 5 min. By 120 min, this perivascular EB aggregation is markedly exacerbated, indicating a severe functional blockade at the gliovascular interface (Scale bar, 10 µm). (D) Quantitative spatial profiling of the perivascular space. The continuous line graph illustrates the mean physical distance (µm) between the CD31^+^ endothelial boundary and the AQP4^+^ astrocytic endfeet along the vessel axis (shaded regions denote ± S.E.M.). 5xFAD mice exhibit a profound and sustained structural widening of the PAS compared to WT controls across all time points, mechanically correlating with the observed fluid stagnation. (E) Plaque-associated fluid entrapment in 5xFAD mice. Left: High-magnification views (from red boxes in B) demonstrating robust EB aggregation (white arrowheads) structurally colocalizing with plaque deposits, a trapping phenomenon persisting from 5 min to 120 min (Scale bar, 5 µm). Right: Representative wide-field confocal images of cortical layer 6 (Ctx L6) co-labeled for EB (green) and ApoE (red). These images illustrate the widespread distribution of plaque-associated CSF entrapment across the deep cortical parenchyma at corresponding time points (Scale bar, 100 µm). (F) Quantitative analysis of plaque-associated EB entrapment in cortical layer 6. Left: Bar graph showing the number of EB+ plaque regions at 5 min and 120 min in 5xFAD mice, indicating a statistically significant decrease over time (*P* < 0.05). Right: Violin plots revealing a concurrent, statistically significant reduction in the cross-sectional area (µm²) of individual EB+ plaques at 120 min compared to 5 min (*P* < 0.05). (G) Magnified views (from solid white boxes in b) displaying the temporal progression of EB recirculation into CD31^+^ vascular lumens. At 5 min, the EB signal remains largely confined to the perivascular space with minimal intraluminal detection. By 120 min, distinct, compartmentalized EB signals (white arrowheads) are robustly detected inside the CD31^+^ vessels in WT mice, representing delayed vascular return. In contrast, this intraluminal re-entry is visibly diminished in the 5xFAD group (Scale bar, 5 µm). (H) Quantitative bar graph of the mean EB fluorescence intensity inside CD31^+^ vessels. In WT mice, intraluminal EB intensity significantly increases from 5 min to 120 min, confirming active vascular recirculation. Crucially, at 120 min, the 5xFAD group exhibits a significant reduction in intraluminal EB intensity compared to WT (*P < 0.05, ***P < 0.001), demonstrating a fundamental failure of the delayed vascular return loop in AD. Bar graph data are presented as mean ± S.E.M. with individual data points.

This profound fluid stagnation physically forced the gliovascular interface apart. Quantitative spatial profiling confirmed that while WT mice maintained a tightly apposed CD31–AQP4 architecture with a separation of less than 1 µm, 5xFAD mice suffered a severe gliovascular decoupling, widening the PAS distance to approximately 4 µm (Figure 3D). This pathological structural expansion acts as a formidable physical barrier, mechanically hindering CSF from effectively entering the deep parenchyma. Furthermore, this spatial decoupling and the loss of intimate AQP4 perivascular apposition strongly corroborate recent findings that the mislocalization of AQP4 directly impairs glymphatic exchange and drives severe brain-wide interstitial fluid stagnation (20, 21).

Furthermore, the limited fraction of CSF that managed to bypass this perivascular bottleneck and diffuse into the tissue was subsequently sequestered by amyloid plaques (Figure 3B and 3E). At 5 minutes post-injection, robust EB signals strongly colocalized with amyloid deposits in the 5xFAD cortex. To comprehensively quantify this entrapment, we analyzed the deeper cortical layer 6, which harbors a high plaque burden. Strikingly, both the absolute number of EB-positive plaques and the cross-sectional area of EB tracer entrapped within these plaques were significantly reduced by 120 minutes compared to the 5-minute baseline (Figure 3F). These temporal kinetics suggest that amyloid plaques function as active physical sinks—initially capturing a large volume of the infiltrating CSF and only slowly releasing it over time—thereby further retarding fluid transit through the pathological parenchyma. This direct physical sequestration of CSF by amyloid deposits provides a critical mechanistic link supporting recent observations that early suppression of glymphatic clearance not only precedes but actively exacerbates amyloid plaque formation, establishing a pathological feed-forward loop of fluid stagnation (22, 23).

Finally, we evaluated the ultimate fate of parenchymal fluid by examining vascular re-entry. Canonical models posit that CSF/ISF primarily exits the parenchyma via venous efflux routes. However, our observations reveal a distinct arterial recirculation pathway. At 5 minutes post-injection, the CD31^+^ arterial lumens were largely devoid of EB signal. Strikingly, by 120 minutes, pronounced, compartmentalized EB signals were detected inside the CD31^+^ arterial lumens of WT mice, visually confirming active delayed vascular return (Figure 3B and 3G). This compartmentalized intraluminal accumulation strongly suggests that fluid re-entry is not a passive diffusive process, but rather relies on the active transcytosis driven by the highly specialized vesicular transport and exosome-mediated machinery intrinsic to brain microvascular endothelial cells (24, 25). In the 5xFAD model, this intraluminal recirculation was severely blunted. Quantitative analysis demonstrated that the mean EB fluorescence intensity within CD31^+^ vessels was significantly reduced in 5xFAD mice at 120 minutes compared to WT controls (Figure 3H).

Taken together, these data demonstrate that the brain-wide collapse of CSF flow in AD is driven by a cascading series of pathological bottlenecks. Upstream macroscopic constriction at the ventral base is compounded by severe microscopic disruptions in the dorsal cortex, including perivascular stagnation, gliovascular decoupling, and parenchymal plaque entrapment. Ultimately, these combined structural and functional barriers culminate in the fundamental failure of the delayed arterial recirculation loop.

### Pathological lipid retention in pericytes and neurons obstructs capillary clearance

To elucidate the cellular mechanisms underlying the failure of macroscopic fluid circulation, we investigated the parenchymal lipid metabolic response following the infiltration of exogenous CSF tracers. As summarized in our working model (Figure 4A), the brain actively mounts a clearance response to remove exogenous materials introduced via I.C.M. injection. In physiological states, this clearance process triggers a rapid, brain-wide elevation in parenchymal cholesterol levels. However, in the AD brain, this initial cholesterol mobilization is significantly blunted.

**Figure 4.**
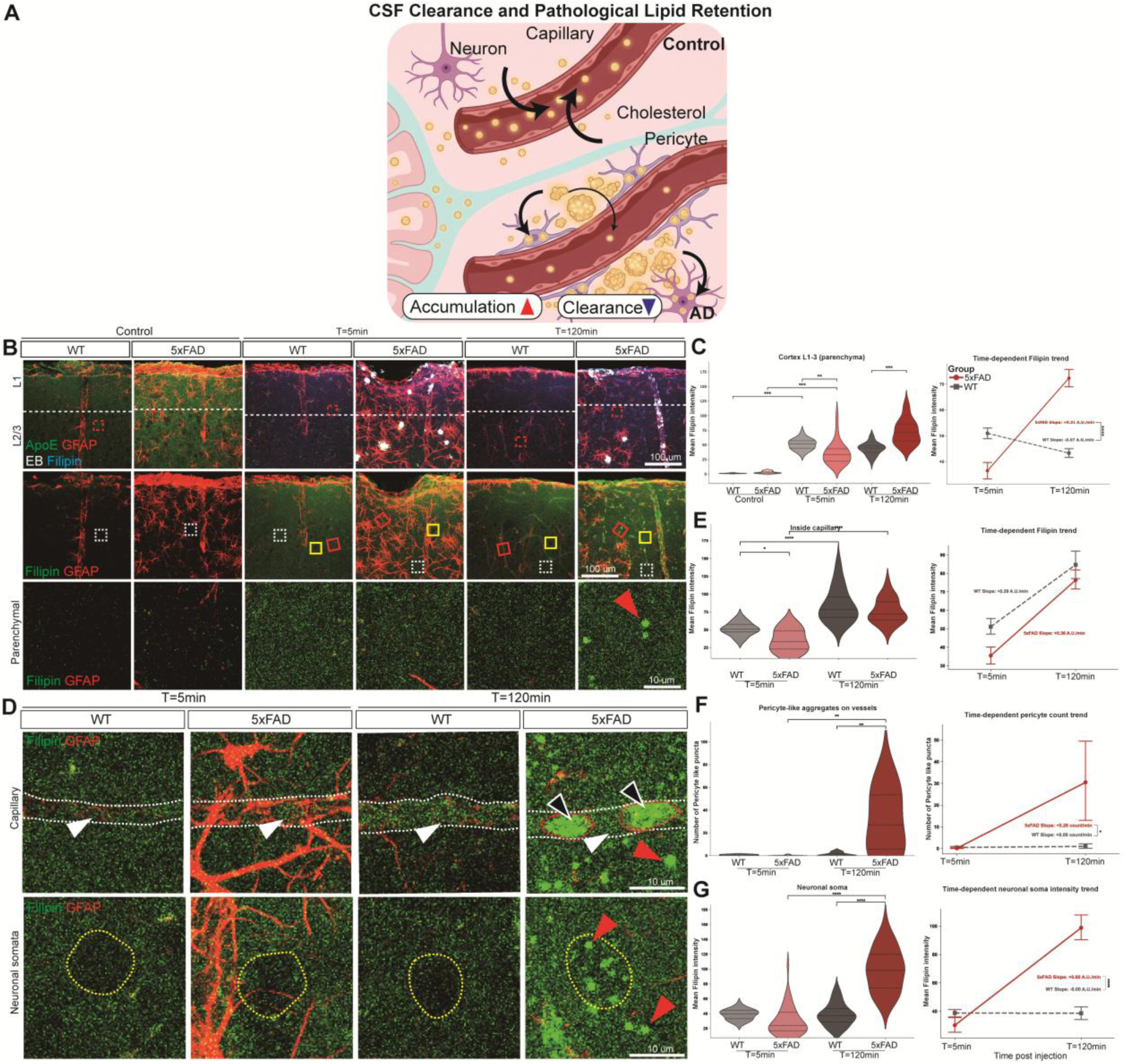
Pathological lipid retention in pericytes and neurons obstructs capillary clearance in Alzheimer’s disease. **(A)** Schematic illustration summarizing the proposed mechanism of cerebrospinal fluid (CSF) clearance and pathological lipid retention. Following the influx of exogenous CSF tracers, parenchymal cholesterol synthesis is rapidly upregulated to facilitate clearance. In wild-type (WT) brains, this mobilized cholesterol is efficiently directed toward and cleared through the capillaries by 120 min. Conversely, in Alzheimer’s disease (AD) models, the initial cholesterol response is blunted, and subsequent capillary efflux is severely impaired. Consequently, uncleared cholesterol retrogradely accumulates, forming massive aggregates within perivascular pericytes and neuronal somata. **(B)** Representative confocal micrographs of the cerebral cortex (layers L1–3) in WT and 5xFAD mice at baseline (Control), 5 min, and 120 min post-injection. Top: Merged images labeled for ApoE (green), GFAP (red), Evans Blue (white), and Filipin (cyan/blue). Middle: Separated channels for Filipin and GFAP. Bottom: High-magnification views of the parenchymal regions corresponding to the white dashed boxes in the middle panels. Scale bars: 100 µm (top, middle); 10 µm (bottom). **(C)** Quantitative spatial profiling of parenchymal Filipin intensity. Left: Violin plots illustrating the mean Filipin intensity distributions across groups. Right: Line graph demonstrating the time-dependent trajectory of parenchymal cholesterol mobilization from 5 min to 120 min. **(D)** High-magnification confocal images of capillary (top, corresponding to red solid boxes in b) and neuronal soma (bottom, corresponding to yellow solid boxes in b) regions (Scale bars, 10 µm). Top: Capillary boundaries are delineated by white dashed lines. White arrowheads indicate intraluminal Filipin signals. Strikingly, in 5xFAD mice at 120 min, distinct pericyte-like morphologies harboring dense lipid accumulations are visible (red dashed lines and black arrowheads), alongside small parenchymal Filipin puncta (red arrowheads). Bottom: Neuronal somata are outlined by yellow dashed lines. By 120 min, 5xFAD mice exhibit pronounced intracellular Filipin aggregation and distinct lipid puncta (red arrowheads) within the neurons, a pathological retention absent in WT and early-stage 5xFAD groups. **(E)** Quantification of intraluminal capillary Filipin intensity. Left: Violin plots showing that at 5 min, the 5xFAD group exhibits significantly reduced intraluminal cholesterol compared to WT. While both groups show a significant increase by 120 min, the time-dependent trend lines (right) reveal no significant difference in their overall slopes, likely confounded by the massive perivascular accumulation in the AD model. **(F)** Quantitative analysis of vessel-associated pericyte-like puncta. Left: Violin plots demonstrating a massive, statistically significant surge in pericyte-like aggregates specifically in the 5xFAD group at 120 min. Right: Trend line graph confirming a significantly divergent accumulation trajectory in AD compared to the flat WT baseline. **(G)** Quantification of Filipin intensity within neuronal somata. Left: Violin plots revealing a significant, delayed intracellular lipid retention in 5xFAD neurons at 120 min, whereas WT levels remain stable. Right: Trend line graph illustrating a significantly accelerated neuronal lipidosis trajectory in the AD model (*P < 0.05, P < 0.01, ***P < 0.001, ****P < 0.0001).

Over a 120-minute period, the mobilized cholesterol in healthy brains is efficiently directed toward the capillaries for vascular efflux, subsequently restoring parenchymal lipid homeostasis. Conversely, this vascular clearance is severely impaired in AD. The failure to extrude cholesterol into the microvasculature leads to massive parenchymal lipid retention, forcing the uncleared cholesterol to aggregate and retrogradely accumulate within perivascular pericytes and adjacent neurons. Collectively, these dual deficits—the initial failure to sufficiently upregulate cholesterol synthesis and the subsequent breakdown of capillary efflux—constitute the pathological hallmarks of CSF clearance dysfunction in AD, culminating in severe cellular lipidosis.

To validate this, we performed high-resolution confocal imaging of Filipin (a cholesterol marker) in the cerebral cortex (layers 1–3). Following I.C.M. injection of the EB tracer, both WT and 5xFAD mice exhibited an acute elevation in Filipin fluorescence at 5 minutes post-injection. This initial cholesterol surge was specifically localized to the parenchymal ECM, distinct from the vascular lumens or neuronal somata. However, the magnitude of this response was fundamentally dysregulated in the AD model; 5xFAD mice displayed significantly lower parenchymal cholesterol levels compared to their WT counterparts at this early 5-minute time point (Figure 4B and 4C). Over the 120-minute clearance period, the two groups demonstrated diametrically opposed lipid trajectories. In WT mice, parenchymal Filipin intensity exhibited a downward trend from 5 to 120 minutes, reflecting the successful clearance of mobilized cholesterol from the tissue. Conversely, the 5xFAD group showed a severe, statistically significant increase in parenchymal cholesterol by 120 minutes. Quantitative trend analysis confirmed that the temporal trajectories of Filipin intensity from 5 to 120 minutes are entirely inverted between the WT and 5xFAD groups, yielding a statistically significant difference in their slopes (Figure 4B and 4C).

Together, these dynamic temporal shifts suggest that a rapid and robust generation of cholesterol is a prerequisite for effectively driving CSF clearance. In AD, this cholesterol-mediated clearance machinery is intrinsically blunted and temporally delayed, ultimately resulting in aberrant lipid accumulation within the brain parenchyma.

The accumulation and dynamic changes of cholesterol during CSF clearance exhibited distinctly different patterns between the WT and 5xFAD groups. By 120 minutes, imaging revealed a general progression of the cholesterol signal toward the vasculature, particularly converging on the capillary networks (Figure 4B). Quantitative analysis of the capillary regions demonstrated that both WT and 5xFAD mice experienced a statistically significant increase in Filipin intensity from 5 to 120 minutes (Figure 4D and 4E). However, in the 5xFAD model, this apparent intraluminal increase is likely a confounding artifact driven by external pathological accumulation. Specifically, high-magnification images of the 5xFAD cortex at 120 minutes revealed massive cholesterol aggregates conforming to pericyte-like cellular structures wrapped directly around the capillaries (Figure 4D, red dashed lines and black arrowheads), which, combined with the overall elevation in surrounding parenchymal cholesterol, artifactually amplifies the capillary signal.

To isolate this, we quantified these vessel-associated pericyte-like cholesterol aggregates, confirming they are a pathological hallmark specific to the 120-minute time point in the AD model. These aggregates were virtually absent in WT mice at all time points and in 5xFAD mice at the early 5-minute stage. Trend analysis further highlighted this stark divergence; while the WT trajectory remained perfectly flat with a slope of zero, the 5xFAD group displayed a sharp positive slope, indicating a significant and rapid surge in perivascular accumulation (Figure 4D and 4F).

Beyond the perivascular accumulation at pericytes, Ctx L 2–3 neurons in the AD model exhibited a distinct and alarming pathological signature. In the WT group, neuronal somata remained largely devoid of anomalous cholesterol signals at both the 5-minute and 120-minute time points. Similarly, at the early 5-minute stage, the 5xFAD mice displayed baseline somatic cholesterol levels indistinguishable from those of WT controls. However, by 120 minutes, the 5xFAD group demonstrated a statistically significant surge in intracellular Filipin intensity. Strikingly, this severe neuronal lipid retention manifested as distinct 1–2 µm cholesterol puncta accumulating directly within the somatic compartment (Figure 4D and 4G, yellow dashed outlines and red arrowheads).

Taken together, these results indicate that the impairment of macroscopic CSF flow in AD is intrinsically tied to a catastrophic failure of the cholesterol-mediated clearance machinery. The inability to effectively extrude mobilized cholesterol into the vasculature leads to massive parenchymal lipid stagnation. Consequently, this unresolvable lipid burden drives the pathological retrograde accumulation of cholesterol into both perivascular pericytes and vulnerable neurons. This terminal cellular lipidosis likely constitutes a core pathogenic mechanism underlying the neuronal toxicity and tissue damage secondary to CSF clearance collapse in dementia.

## Discussion

Our findings establish a comprehensive brain-wide CSF recirculation loop that enters via the PAS of major ventral arteries—namely the CW, OlfA, and MCA—and propagates toward dorsal regions. Macroscopic imaging reveals that in AD mouse model, structural abnormalities within this ventral PAS network precipitate a severe stagnation of CSF flow, characterized by reduced dorsal entry and abnormal, asymmetric ventral diffusion (Supplementary Figure 1, and Supplementary Figure 2). This indicates that the macroscopic collapse of dorsal CSF clearance in dementia is fundamentally driven by upstream structural bottlenecks at the ventral arterial roots.

The severity of these ventral structural defects is visually and quantitatively evident (Figure 2). IVIM imaging—which we specifically optimized for structural tracking up to a functional maximum depth of 200 µm—demonstrates a pronounced narrowing of both the MCA lumen and its surrounding PAS in the 5xFAD group. Although a ∼30% reduction in the overall MCA-PAS diameter (Figure 2E) may appear structurally modest, its hydrodynamic consequence could be disproportionately large. To illustrate this, we approximated the PAS as an annular channel bounded by the vascular wall and the AQP4 endfoot layer (Figure 2F) and applied Poiseuille-type scaling, in which volumetric flow rate (*Q*) scales approximately with the fourth power of the effective channel radius under the simplifying assumptions of laminar flow and unchanged pressure gradient and fluid viscosity between genotypes (26):

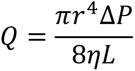

We emphasize that this classical relationship, derived for flow through a simple cylindrical pipe, provides only an order-of-magnitude approximation for the annular, and possibly non-uniform, PAS geometry observed here; precise hydrodynamic modeling would require computational fluid dynamics incorporating the measured inner and outer boundaries. With this caveat, applying the observed ∼15– 30% reduction in effective PAS width (Figure 2F) to this scaling relationship suggests a substantial, potentially several-fold, reduction in periarterial influx capacity, indicating that even modest structural narrowing could translate into a disproportionately large functional bottleneck. Furthermore, we identified highly organized, linear tracks within the PAS during initial influx. These organized tracks suggest that CSF does not flow as a uniform sheet, but may propagate along distinct anatomical micro-channels, potentially aligning with the structural gaps between smooth muscle cell (SMC) filaments or representing entirely novel periarterial tubular architectures; resolving the physical basis of these tracks will require higher-resolution structural imaging.

Recent models argue that passive diffusion alone cannot theoretically account for the comprehensive transit of CSF from the arterial PAS, through the dense parenchyma, and into venous or lymphatic efflux routes (1, 9). Our detergent-free LD and EB imaging provides the first definitive experimental evidence of an active, vesicle-mediated CSF pathway operating across the brain parenchyma (Supplementary Figure 3A). We identified distinct CD31/AQP4/EB-positive vesicular structures co-packaged with albumin and mIgG, confirming that luminal contents do not merely diffuse but are actively transcytosed. Crucially, the size of these transport vesicles is significantly enlarged in the 5xFAD model, a defect that exacerbates by 120 minutes post-injection (Supplementary Figure 3B and Supplementary Figure 4). This suggests that AD pathology disrupts the precise vesicle biogenesis processes orchestrated by brain endothelial cells and astrocytes, resulting in bloated transport complexes that fail to efficiently shuttle clearance cargo to neuronal targets.

The formation of amyloid plaques is a defining hallmark of AD, and our data highlight their highly lipophilic physical nature as a massive physical trap for fluid flow (27, 28). Even prior to tracer injection, LD staining reveals plaque-like lipid aggregates in the 5xFAD cortex (Supplementary Figure 5). Following I.C.M. injection, WT brains remain free of such structures, but 5xFAD mice exhibit robust colocalization of EB and LDs directly at the amyloid plaques. This sequestration indicates that the lipid-dependent CSF transport machinery is physically entangled by the extreme lipophilicity of amyloid deposits. Furthermore, baseline LDs are distributed throughout neuronal somata and the ECM; at 5 minutes post-injection, the 5xFAD group shows an enlargement of these LDs alongside a reduction in total number and ECM signal. By 120 minutes, aberrant LD accumulation surges at the capillary beds. This trajectory suggests that lipid-mediated CSF clearance naturally drains toward capillaries, but in AD, defective vascular efflux traps the lipids, leading to profound parenchymal accumulation.

Canonical glymphatic frameworks largely model a unidirectional flow: from periarterial influx into the parenchyma, culminating in paravenous efflux (3, 29). However, our dynamic tracking reveals a distinct bi-directionality in physiological states, providing direct imaging evidence of arterial recirculation, in which parenchymal fluid re-enters the structural boundaries of arteries by 120 minutes post-injection. In AD, this recirculation appears severely compromised, coinciding with profound gliovascular decoupling in the dorsal cortex (Supplementary Figure 6). The pathological accumulation of tracer forms massive aggregates within the PAS as early as 5 minutes post-injection, physically widening the distance between the CD31⁺ endothelial layer and the AQP4⁺ astrocytic endfeet. To estimate the plausible biophysical consequence of this widening, we modeled diffusive exchange across the perivascular gap using Fick’s First Law, where the flux (*J*) of solutes is proportional to the concentration gradient across that distance (*dx*) (30):

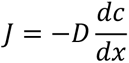

and the characteristic diffusion time (*t*) scales with the square of the diffusion distance:

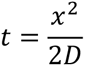

Assuming a comparable concentration gradient and diffusion coefficient between genotypes—an approximation, since neither was measured directly in this study—the observed ∼4-fold expansion of the gliovascular gap (from <1 µm in WT to ∼4 µm in 5xFAD) would be expected to correspond to an approximately 4-fold reduction in diffusive flux and a 16-fold increase in diffusion time. While these estimates should be interpreted as order-of-magnitude approximations rather than measured flow rates, they illustrate how even a modest structural expansion of the gliovascular interface could translate into a substantial kinetic barrier to solute exchange, consistent with the perivascular fluid stagnation observed in the AD cortex. Whether this structural widening is itself a primary cause of impaired exchange, or instead a secondary consequence of upstream vesicular trafficking failure that causes fluid to back up and mechanically distend the PAS, cannot be resolved by the present time-course data and will require targeted interventional studies to disambiguate.

Astrocytes play an indispensable role in CNS lipid homeostasis, orchestrating cholesterol synthesis and ApoE-mediated lipid assembly (31, 32). Pre-injection imaging displays widespread parenchymal ApoE expression; however, following CSF tracer influx, WT brains show ApoE highly concentrated within astrocytes to drive lipid clearance. In stark contrast, 5xFAD brains fail to mount this targeted astrocytic response. Instead, ApoE remains diffusely spread and heavily concentrated on amyloid plaques—a distribution completely absent in the un-injected state (Supplementary Figure 7 and Supplementary Figure 8). Concurrently, total cholesterol levels surge dramatically following injection, underscoring that brain-wide fluid clearance is fundamentally a lipid-driven process. We hypothesize that in AD, reactive astrocyte states disable this acute lipid clearance machinery, leaving functional ApoE trapped in pathological aggregates.

Furthermore, WT mice successfully extrude cholesterol and EB into the capillary beds by 120 minutes, revealing an active *arterial* recirculation system that challenges traditional venous-only efflux models. In 5xFAD mice, this capillary clearance is blocked, resulting in pericyte-like structures abnormally laden with cholesterol along the arteries (Supplementary Figure 7 and Supplementary Figure 8). This identifies pericytes as essential mediators of lipid efflux into the vasculature. In AD, pericytes likely undergo a pathological state shift, losing their ability to express key lipid efflux transporters like ABCA1 (which normally mediates ApoE-dependent cholesterol efflux from brain pericytes), thereby transforming from functional clearance gateways into pathological lipid sinks (33, 34).

Amyloid plaques exert their influence not only through chemical toxicity but also through complex biophysical interactions (35, 36). High-resolution Correlative Light and Electron Microscopy (CLEM) and TEM analyses demonstrate that plaques are not mere fibril aggregates (Supplementary Figure. 9 and Supplementary Figure 10). Instead, they are highly heterogeneous structures containing pre-and post-synaptic debris, myelin fragments, and remarkably small lipid vesicles trapped within massive membrane architectures. We also identified cell-like structures with exceptionally high lipid content within these plaque regions (Supplementary Figure 9). While these baseline un-injected structures illustrate the lipophilic nature of the AD microenvironment, future CLEM studies tracking post-injection tracer dynamics will be essential to map exactly how these pathological hubs entangle and dismantle incoming CSF transport vesicles.

To successfully permeate the brain parenchyma, luminal fluids must first bypass the endothelial barrier. In our in vitro Transwell assays utilizing HBMVECs, control groups (dPBS) showed no crossover of albumin, whereas the application of 5% BSA to the upper chamber resulted in substantial albumin transport to the lower compartment (Supplementary Figure 11). This active, directional shuttling confirms that brain endothelial cells possess the intrinsic capability to transcytose bulk macromolecular cargo, highly supporting the existence of a physiological albumin-based vesicular shuttling system bridging the vasculature and the deep brain matrix.

Building upon this endothelial mechanism, we investigated whether the albumin-vesicle pathway could be exploited systemically. Following retro-orbital injection, the administration of an endothelial-processed albumin and EB mixture dramatically amplified the abundance of parenchymally infiltrated CD31^+^/EB^+^ vesicles at 30 and 60 minutes compared to EB alone (Supplementary Figure 12). This dynamic presents a groundbreaking therapeutic strategy. Because the BBB stringently restricts the entry of most conventional neurodegenerative drugs, packaging therapeutics within albumin-binding conjugates or lipid nanocarriers designed to hijack this endogenous endothelial transcytosis route could finally bypass the BBB, enabling targeted drug delivery deep into the pathological parenchyma (37, 38).

In our imaging assays, baseline 647 nm excitation in un-injected control brains produced weak, yet structurally defined, vascular and vesicular signals rather than random optical noise. This baseline phenomenon likely originates from the weak intrinsic autofluorescence or optical scattering arising from the complex structural conformations of densely packed endogenous proteins, such as circulating albumin, within the vascular lumen (39). More critically, while physiological CSF and artificial CSF (aCSF) lack albumin, our in vitro microplate assays revealed a profound chemical synergy: whereas EB or BSA alone in aCSF yields a minimal 8-fold baseline fluorescence, the binding of EB to BSA results in an exponential, >4,300-fold amplification in emission intensity under 647 nm excitation (Supplementary Figure 13). This synergistic amplification dictates that the intensely bright EB signals observed in recirculating vesicles and the parenchyma represent structurally intact, protein-bound complexes rather than free diffusion. Not all tissue EB colocalizes with albumin, suggesting that these intense signals also arise from EB strongly binding to other abundant parenchymal lipid or cholesterol carriers.

This study delineates the multifaceted collapse of CSF recirculation in dementia; however, several critical mechanisms warrant future investigation. 1) Ventral Architecture: The precise ultrastructural identity of the linear CSF tracks within the ventral arterial PAS remains to be defined via correlative electron microscopy. 2) Transcellular Routing: The exact molecular mechanisms by which CD31^+^ transport vesicles traverse the blood-brain barrier, navigate the extracellular matrix, deliver cargo to neurons, and recirculate into the arteries require detailed multi-omics tracing. 3) Metabolic Triggers: The signaling cascades that allow the brain to instantly detect CSF infiltration and trigger acute, brain-wide cholesterol synthesis for fluid clearance must be elucidated to fully understand this endogenous defense system.

## Methods

All experimental protocols and analytical methodologies are detailed in the Supplemental Methods.

### Sex as a biological variable

This study utilized both male and female mice, as the mechanisms of CSF recirculation and AD pathogenesis investigated here are expected to be relevant to both sexes.

### Statistics

All statistical and comparative analyses were performed programmatically using the scipy.stats(40) and statsmodels(41) libraries in Python. Quantitative data were assessed for statistical significance using unpaired two-tailed Student’s t-tests for two-group comparisons, or ANOVA followed by Tukey’s honestly significant difference (HSD) post-hoc tests to correct for multiple comparisons across three or more experimental groups (e.g., WT vs. 5xFAD; control vs. time-course treatments). For kinetic influx data, two-way ANOVA was employed to evaluate interaction effects (genotype × time). All graphical data were generated using seaborn and are presented as mean ± S.E.M. P-values < 0.05 were considered statistically significant (* P < 0.05, P < 0.01, *** P < 0.001, **** P < 0.0001).

### Study approval

All animal experiments were performed in accordance with the guidelines and protocols approved by the Institutional Animal Care and Use Committee (IACUC) of the Korea Brain Research Institute (KBRI) (Approval No: IACUC-26-00026).

### Data availability

The datasets generated and analyzed in this study—including raw multidimensional imaging data, image processing outputs, microplate reader results, and detailed records of mouse model production and tissue sectioning—have not been deposited in a public repository. However, all raw data and supporting materials are readily available from the corresponding author upon reasonable request.

## Author Contributions

M.C., and D-G.K. conceived and designed the study. M.C., K-S.K., I-W.K., and M.K.J. developed the methodology and analysis pipeline. M.C., K-S.K., I-W.K., and M.K.J. performed the investigations, including animal experiments, and imaging. M.C., K-S.K., and I-W.K. carried out the formal data analysis, encompassing bioinformatics, statistical evaluation, and modeling. D-G.K. provided resources and secured funding. M.C. wrote the original draft of the manuscript. All authors (M.C., K-S.K., I-W.K., M.K.J., and D-G.K.) contributed to reviewing and editing the manuscript. Visualization of figures and data plots was performed by M.C. Supervision was provided by M.C., and D-G.K. Project administration was managed by D-G.K.

## AI Disclosure

Google Antigravity (Google LLC) was utilized between January 2026 and August 2026 to assist in manuscript writing, developing custom Python scripts for computational analyses (detailed in Supplemental Methods), and generating the results summary illustration for the main figure. The authors assume all responsibility for ensuring the integrity and accuracy of the AI output presented in this manuscript.

## Supplementary material

Supplemental data (Methods and Figures (13))

## Acknowledgements

We thank members of the Brain Research Institute for technical support and constructive discussions.

Address correspondence to:

Min Kyo Jung, Neural Circuit Research Group, Korea Brain Research Institute, Daegu, Republic of Korea. Phone: 82.53.9808424 ;

Do-Geun Kim, Dementia Research Group, Korea Brain Research Institute, Daegu, Republic of Korea. Phone: 82.53.9808364 ;

