## Supplementary Figures for "Brain-wide Cerebrospinal Fluid Recirculation Collapse in Alzheimer’s Disease"

Supplemental Methods

### Animals

All animal experiments were performed in accordance with the guidelines approved by the Institutional Animal Care and Use Committee (IACUC) of the Korea Brain Research Institute (KBRI). Adult male and female 5xFAD TG and their littermates as WT (6-month-old), B6129 (8–12 weeks old) were used in this study. Animals were housed under a controlled 12-hour light/dark cycle with ad libitum access to food and water. (IACUC-26-00026/KBRI)

### Tracer Preparation and CM Injection

Mice were anesthetized via an intraperitoneal injection of Avertin (2.5% tribromoethanol, 0.02 mL/g). To ensure precise and stable delivery of tracers into the CSF compartment, a custom-made cannula system was prepared based on a previously established cisterna magna cannulation protocol (1), with minor modifications. This system consisted of a Hamilton syringe connected to polyethylene (PE) tubing (SP-10; inside diameter, 0.28 mm; outside diameter, 0.61 mm), with a 30-gauge needle tip attached to the distal end to serve as the injection port. Under deep anesthesia, the skin and muscle layers overlying the posterior neck were surgically retracted to expose the Atlanto-occipital membrane, providing direct access to the CM. The 30-gauge needle tip of the pre-filled cannula was carefully inserted into the CM and rigidly secured to the surrounding tissue and skull using dental cement to prevent leakage of CSF or tracer and ensure structural stability. Following fixation, 5 µl of 4% EB was infused at a constant rate of 1 µl/min. To minimize backflow and maintain stable intracranial pressure, the cannula remained in place for at least 5 minutes post-infusion. The animals were subsequently sacrificed at specified time points (5, 15, 30, 60, and 120 min) for downstream analysis.

### In Vitro HBMVEC Culture

BBB Permeability Assay: HBMVECs obtained from Cell Systems were cultured in complete classic medium at 37°C in a humidified atmosphere with 5% CO2. To establish an BBB model, transwell permeable inserts (3.0 µm pore size) were pre-coated with attachment factor (Cell Systems, Cat# 4Z0-210) according to the manufacturer's instructions. HBMVECs were seeded onto the apical side of the coated inserts, and culture media were refreshed every 3 days. The cells were allowed to grow and differentiate for 7 days post-seeding to establish a fully confluent and tight endothelial monolayer prior to conducting transcytosis assays. After 7 days of cultivation to establish a fully confluent endothelial monolayer, the basolateral medium was replaced with pre-warmed Hanks' Balanced Salt Solution (HBSS), while complete culture medium was retained in the apical chamber. A 40 % BSA stock solution was added into the apical chamber to achieve a final working concentration of 5% BSA. Following incubation at 37°C for 1 h, the entire solution from the basolateral chamber was collected. The concentration of transported albumin was quantified by mixing the collected basolateral samples with EB solution and measuring the absorbance at 620 nm using a microplate reader.

**Tissue Processing and Eci Clearing**

Following specified circulation times post-CM injection, mice were anesthetized with Avertin (2.5% tribromoethanol, 0.02 ml/g). Brains were rapidly harvested without cardiac perfusion to prevent the washout of tracers and preserve their endogenous distribution for macroscopic analysis. Whole brains were immediately subjected to optical photography and fluorescence mapping using a ChemiDoc imaging system to visualize the surface distribution of EB-albumin complexes. Subsequently, brains were divided into hemibrains for distinct downstream applications: one hemibrain was fixed in 4% PFA at 4°C overnight for tissue clearing and light-sheet microscopy, while the contralateral hemibrain was fixed in 2% PFA for sectioning and immunofluorescence staining. The 4% PFA-fixed hemibrains were washed with 1xPBS three times (30 minutes each) on a shaker at room temperature to remove residual fixative. Tissue clearing was performed using a streamlined Ethyl Cinnamate (ECi)-based protocol, as previously described (2), with minor modifications. Dehydration was achieved through a graded ethanol series (50% for 1 hour, 70% for 1 hour, 100% for 1 hour twice, and 100% overnight). Dehydrated tissues were immersed in undiluted ECi (Sigma-Aldrich, Cat# W243000) and incubated overnight at room temperature with gentle shaking, strictly protected from light until the tissue achieved amber-like transparency. Cleared hemibrains were imaged using a LSFM with a refractive index (RI) matching of 1.56. Raw image data were converted into IMARIS files, stitched, and processed for 3D reconstruction and maximum intensity projection. Quantitative analysis of the glymphatic distribution and tracer volumes was performed using the reconstructed 3D volumes.

**Immunofluorescence and Histology**

Following extraction and postfixation, coronal and sagittal brain sections (30–50 μm in thickness) were prepared using a vibrating blade microtome (vibratome). Free-floating sections were permeabilized with 1% Triton X-100 in phosphate-buffered saline (PBS) for 20–30 minutes at room temperature, followed by blocking with 5% normal goat serum (NGS). Subsequently, the sections were incubated overnight at 4 °C with the following primary antibodies: rat anti-albumin (1:500; Takara Bio, M234), rat anti-CD31 (1:500; BD Biosciences, #550274), mouse anti-glial fibrillary acidic protein (GFAP; 1:500; Cell Signaling Technology, 3670S), mouse anti-aquaporin-4 (AQP4; 1:500; Abcam, ab9512), rabbit anti-apolipoprotein E (ApoE; 1:500; Invitrogen, Thermo Fisher Scientific, 701241), and mouse anti-NeuN (1:500; MilliporeSigma, MAB377).

After extensive washing, the sections were incubated with appropriate Alexa Fluor-conjugated secondary antibodies (1:500; Invitrogen, Thermo Fisher Scientific) for 2 hours at room temperature. The specific secondary antibodies utilized included Alexa Fluor 488-, 568-, or 647-conjugated anti-mouse IgG (A11001, A11004, A21235), anti-rabbit IgG (A11008, A11011, A32733), and anti-rat IgG (A11006, A11077, A21247). For pathological and lipid labeling, fibrillar amyloid plaques were counterstained with Thioflavin S (Sigma-Aldrich, Merck KGaA, T1892). Intracellular lipid droplets and unesterified cholesterol were visualized using LipidSpot™ 488/610 (1:1,000; Biotium, 550274) and Filipin III complex (25 μg/ml; Sigma-Aldrich, Merck KGaA, SAE0087), respectively, according to the manufacturers' instructions.

High-resolution multi-channel fluorescence images were acquired using a STELLARIS 8 confocal laser scanning microscope (Leica Microsystems GmbH). For large-scale regional mapping and whole-brain section scanning, an inverted motorized epifluorescence microscope (Ti-E; Nikon Instruments) was utilized. All imaging parameters (laser power, gain, and pinhole size) were kept constant across comparative groups to ensure quantitative reliability.

To analyze the macroscopic CSF flow and three-dimensional distribution of the EB tracer across the entire brain, light-sheet fluorescence microscopy (LSFM) was performed. Cleared samples were retrieved, and residual ECi on the tissue surface was carefully removed using laboratory wipes. To ensure stable and vertical mounting during imaging, a razor blade was used to trim approximately 2 mm of the cerebellar region, creating a flat, uniform cross-sectional surface. This flattened cerebellar face was adhered using a gel-type superglue onto a custom-designed mounting stand fabricated via 3D printing, allowing the whole brain to stand upright. The mounted tissue was placed into the imaging chamber filled with fresh ECi as the refractive index-matching immersion solution. Whole-brain imaging was performed using a light-sheet fluorescence microscope (Ultramicroscope II; Miltenyi Biotec). Subsequent image processing, including tile stitching and 3D reconstruction, was executed using the IMARIS software suite to generate comprehensive whole-brain volumetric models.

Macroscopic fluid dynamics and tracer distribution were evaluated using an IVIM imaging system. Following C.M. injection and tissue harvesting, the cleared brain tissues were embedded and stabilized in a 60 mm petri dish using a 2% agarose mold filled with ECi for refractive index matching. A coverslip was carefully placed over the tissue, and the mounted sample was secured on the IVIM stage. High-resolution multi-position Z-stack imaging was performed using a 10x objective lens. Image processing, including tiling/montage, 3D reconstruction, and cross-sectional image generation, was executed using the dedicated software provided with the IVIM system to facilitate downstream quantitative analysis.

Correlative Light and Electron Microscopy (CLEM) Following initial observation under a fluorescence microscope, mouse brain slices containing the region of interest were trimmed into 1 mm × 1 mm sections (3). The tissues were fixed in a solution containing 2% paraformaldehyde (EM grade; Electron Microscopy Sciences (EMS), #19210) and 2.5% glutaraldehyde (EMS, #16210) diluted in 0.1 M sodium cacodylate buffer. After thorough washing, the samples were post-fixed in 2% osmium tetroxide (OsO4) supplemented with 1.5% potassium ferrocyanide for 1 hour at 4 °C. The specimens were subsequently dehydrated through a graded acetone series (50%, 60%, 70%, 80%, 90%, and 100% for 20 min at each concentration) and infiltrated with an epoxy resin (EMS, #14120). The samples were flat-embedded between Aclar films (EMS, #50425), polymerized at 60 °C, and mounted on blank resin blocks in preparation for serial sectioning. Ultrathin sections (60 nm) were cut using an ultramicrotome (Leica Microsystems GmbH), and double-stained with UranyLess (EMS, #22409) for 2 min and 3% lead citrate (EMS, #22410) for 1 min. Electron micrographs were acquired using a Tecnai G2 transmission electron microscope (TEM) operated at 120 kV (Thermo Fisher Scientific).

To generate correlative images, fluorescence and corresponding TEM images were overlaid using image editing software (ImageJ and Affinity Photo) (4). The opacity of the fluorescence channel was reduced to 50%, and structural alignment was achieved by precisely adjusting the magnification (preserving the aspect ratio) and rotation angle of the confocal images to match the TEM ultrastructure, utilizing distinctive structural landmarks clearly visible in both modalities.

**Image Processing and Automated Computational Analysis**

To ensure reproducibility, minimize human bias, and strictly quantify morphological and dynamic changes across experimental groups, all image processing and statistical analyses were performed programmatically using custom-built Python algorithms (Python 3.8+, utilizing OpenCV, scikit-image, and SciPy libraries). The computational codes were generated with the assistance of Google Antigravity (Google LLC). The complete analysis pipelines and source codes have been made publicly available on GitHub (https://github.com/KBRIDGLAb/CSF_flow_project).

**In Vitro Permeability and Microplate Assays**

To evaluate *in vitro* barrier integrity, albumin and EB concentrations across HBMVEC porous membrane compartments were fluorometrically quantified. Raw microplate reader datasets were parsed and log-transformed where necessary to normalize the wide dynamic ranges of fluorescence intensity. Statistical differences across interaction conditions (e.g., EB, BSA, aCSF) were evaluated using one-way ANOVA followed by Bonferroni-corrected t-tests.

**Extravascular Vesicle and Puncta Subpopulation Analysis**

To specifically quantify extravascular vesicles and permeated puncta while strictly excluding those confined within or attached to the macro-vascular endothelium, a dual-layer spatial masking and extraction algorithm was developed. First, macro-vascular structures were aggressively masked to define the exclusion zone by subjecting the structural channel (e.g., CD31) to Gaussian blurring(5) and Otsu's thresholding (6). Contiguous components exceeding a stringent maximum size threshold were classified as macro-vasculature and subsequently dilated to completely encapsulate the endothelial border and luminal space, creating a definitive vascular exclusion mask. Within the remaining extravascular space (ECM and neural tissue), vesicular structures were extracted using an elliptical morphological Top-Hat filter to isolate highly localized, discrete puncta from the diffuse background fluorescence. To completely eliminate human bias in defining vesicle size categories, the physical areas ($\mu m^2$) of all detected extravascular puncta were subjected to an unsupervised machine learning algorithm, the Gaussian Mixture Model (GMM)(5). The GMM mathematically deconvoluted the continuous size distribution into distinct Gaussian subpopulations (small vs. large vesicles) and automatically established the optimal decision boundary based on intrinsic data density, allowing for highly rigorous and unbiased subpopulation counting.

**Macroscopic Structural Analysis of the MCA**

To assess the macroscopic distribution of the CSF tracer along the ventral vascular structures, specifically the MCA, Z-stack images acquired from the cleared brain tissues were projected and analyzed. Vessel wall boundaries were mathematically mapped to compute continuous distance-over-length profiles of the perivascular space. To quantify the spatial paravascular accumulation of the tracer, concentric circular masks reflecting the inner and outer diameters of the MCA were computationally generated. Vesicular structures and tracer aggregates within these concentric perivascular domains were then isolated and quantified to accurately assess the extent of transmural and paravascular tracer efflux.

**Blood-Brain Barrier Integrity and Vascular Distance Analysis**

To quantitatively evaluate the perivascular localization and polarization of AQP4 astrocyte endfeet, a custom graphical user interface (GUI) was developed in Python to trace physical vascular boundaries. For individual vascular segments, the abluminal $CD31^+$ endothelial wall and the corresponding $AQP4^+$ astrocyte endfeet boundary were manually annotated as continuous polylines. To compute the detachment distance, the AQP4 polyline was computationally interpolated into 100 evenly distributed spatial coordinates. The minimum Euclidean distance from each AQP4 coordinate to the nearest segment of the CD31 polyline was calculated and converted to physical distances (μm) using objective lens metadata, yielding high-resolution spatial profiles of perivascular detachment across experimental groups.

**Amyloid Plaque Morphological Analysis**

Automated detection and morphological quantification of $EB^{+}$ amyloid plaques were performed using a specialized spatial filtering algorithm. To rigorously exclude non-specific fluorescence saturation artifacts inherently present at the cortical surface, the algorithm explicitly cropped and excluded the top 20% of the image field. In the remaining deep cortical and hippocampal regions, individual plaques were segmented, and their physical sizes (area, $\mu m^2$) were calculated by mapping pixel coordinates to objective lens physical scaling metadata. Colocalization analysis was subsequently performed to evaluate the overlap of EB+ plaques with target markers (e.g., ApoE, lipids).

**Intra-vascular Evans Blue Retention Analysis**

To assess the retention and accumulation of EB dye within the vascular lumen, a semi-automated region of interest (ROI) extraction tool was utilized. $CD31^{+}$ blood vessels were traced, and the algorithm computationally extracted both the mean EB fluorescence intensity and the proportional EB-positive area fraction strictly confined within the defined endothelial boundaries, avoiding signal contamination from the surrounding parenchyma.

**Filipin (Cholesterol/Lipid) Transport Dynamics**

To map the complex spatiotemporal dynamics of cholesterol and lipid transport, Filipin fluorescence intensity was quantified across distinct anatomical brain compartments. For pericyte extraction, an automated computer vision pipeline was engineered to selectively extract pericyte bodies. To distinguish pericytes from background noise and vascular fragments, strict morphological gating parameters were applied, including area constraints and a shape-based solidity threshold ($> 0.45$). For compartmental influx and accumulation, mean Filipin intensities were independently mapped and quantified within the brain parenchyma, internal capillary lumens, and neuronal somata.

**Code availability**

Custom Python scripts used for BBB perivascular distance quantification, extravascular vesicle subpopulation clustering (GMM), amyloid plaque morphometry, pericyte extraction, IVIM paravascular dynamics, and spatiotemporal Filipin tracking are publicly available on GitHub (https://github.com/KBRIDGLab/CSF_flow_project). All computational analyses were executed in Python (v3.14.2). Initial code generation was assisted by Google Antigravity (Google LLC). Key packages utilized for data processing, image analysis, and statistical evaluation include OpenCV-Python (v4.13.0), NumPy (v2.4.0), SciPy (v1.16.3), pandas (v2.3.3), statsmodels (v0.14.6), and seaborn (v0.13.2). The exact thresholds, algorithms, and statistical parameters applied for each analysis are detailed in the Methods and the accompanying repository documentation.

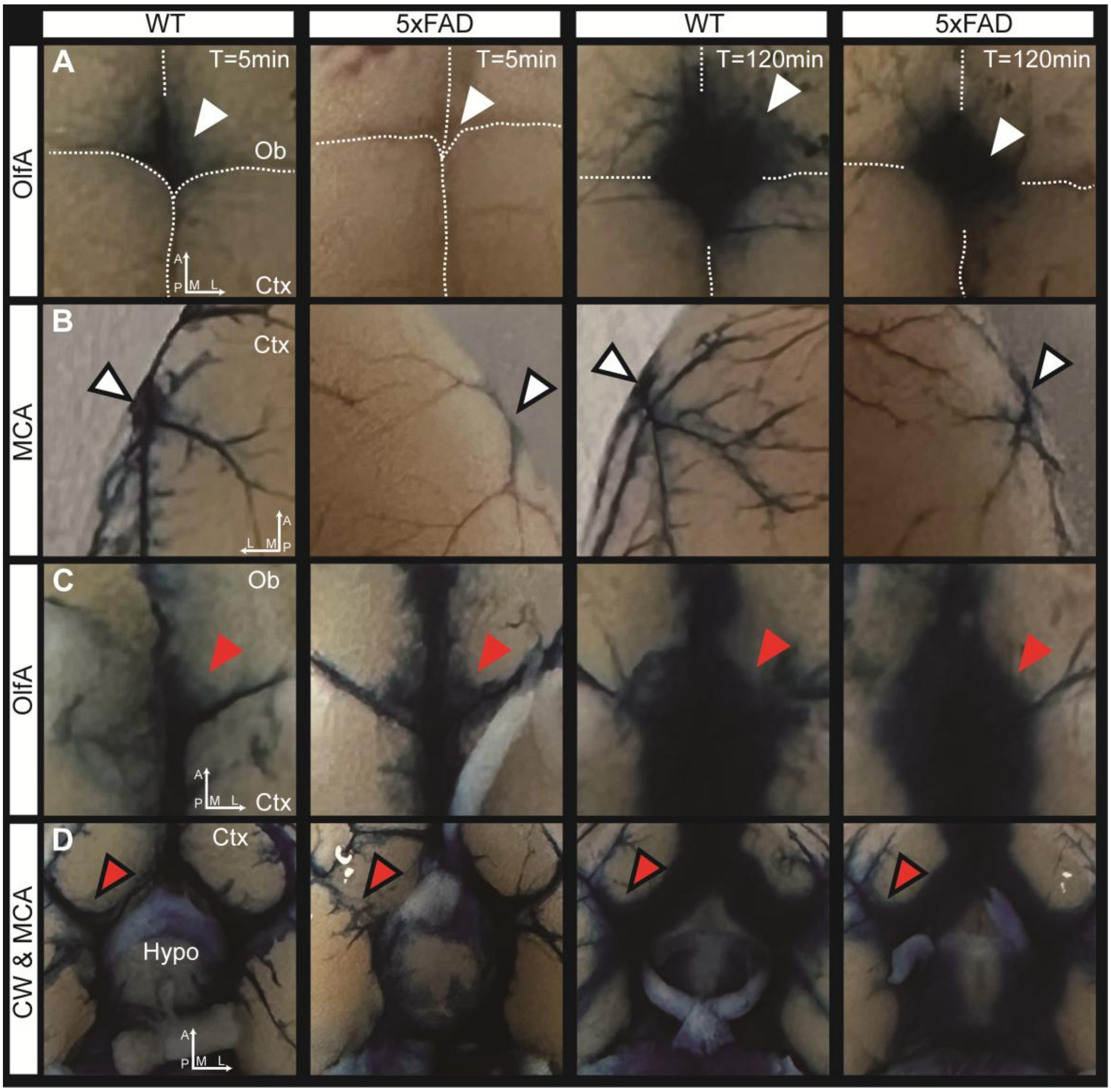


**Supplementary Figure. 1 | Macroscopic flow stagnation and asymmetric distribution of CSF tracer in the ventral and dorsal arterial networks of 5xFAD mice.** Optical zoom images illustrating the distribution of intra-cisterna magna (I.C.M.)-injected Evans Blue (EB) tracer along major arterial tracts—the olfactory artery (OlfA), middle cerebral artery (MCA), and Circle of Willis (CW)—at 5 min and 120 min post-injection in wild-type (WT) and 5xFAD mice. Directional arrows indicate Anterior-Posterior (A-P) and Medial-Lateral (M-L) axes. **(A)** Dorsal view of the OlfA region. The boundaries between the cortex (Ctx) and olfactory bulb (Ob) are delineated by white dashed lines. In the WT brain, the EB tracer robustly propagates along the OlfA into the dorsal territory. Conversely, the 5xFAD group exhibits a marked reduction in dorsal OlfA tracer influx (solid white arrowheads) at both 5 min and 120 min. **(B)** Dorsal view of the MCA region. The MCA trajectory is highlighted by white arrowheads with black outlines. Similar to the OlfA, the dorsal ascent of the EB tracer along the MCA is prominently visible in WT mice but is severely diminished in the 5xFAD model across all time points. **(C)** Ventral view of the OlfA region. Red arrowheads denote the ventral origin of the OlfA. In 5xFAD mice, the tracer fails to efficiently ascend dorsally; instead, it stagnates at the ventral base, leading to an abnormal, broadened diffusion pattern into the surrounding ventral tissues at 120 min compared to the tightly restricted flow observed in WT controls. **(D)** Ventral view encompassing the CW and MCA regions near the hypothalamus (Hypo). Red arrowheads with black outlines indicate the bilateral MCA branches. In WT mice, the tracer distributes evenly and symmetrically into both MCAs. In stark contrast, 5xFAD mice exhibit a severe left-right flow asymmetry as early as 5 min, with tracer propagation drastically blunted on one side. By 120 min, this asymmetric basal pooling results in a widespread, hazy diffusion of the trapped tracer across the ventral surface, further confirming a critical failure in directed arterial influx.

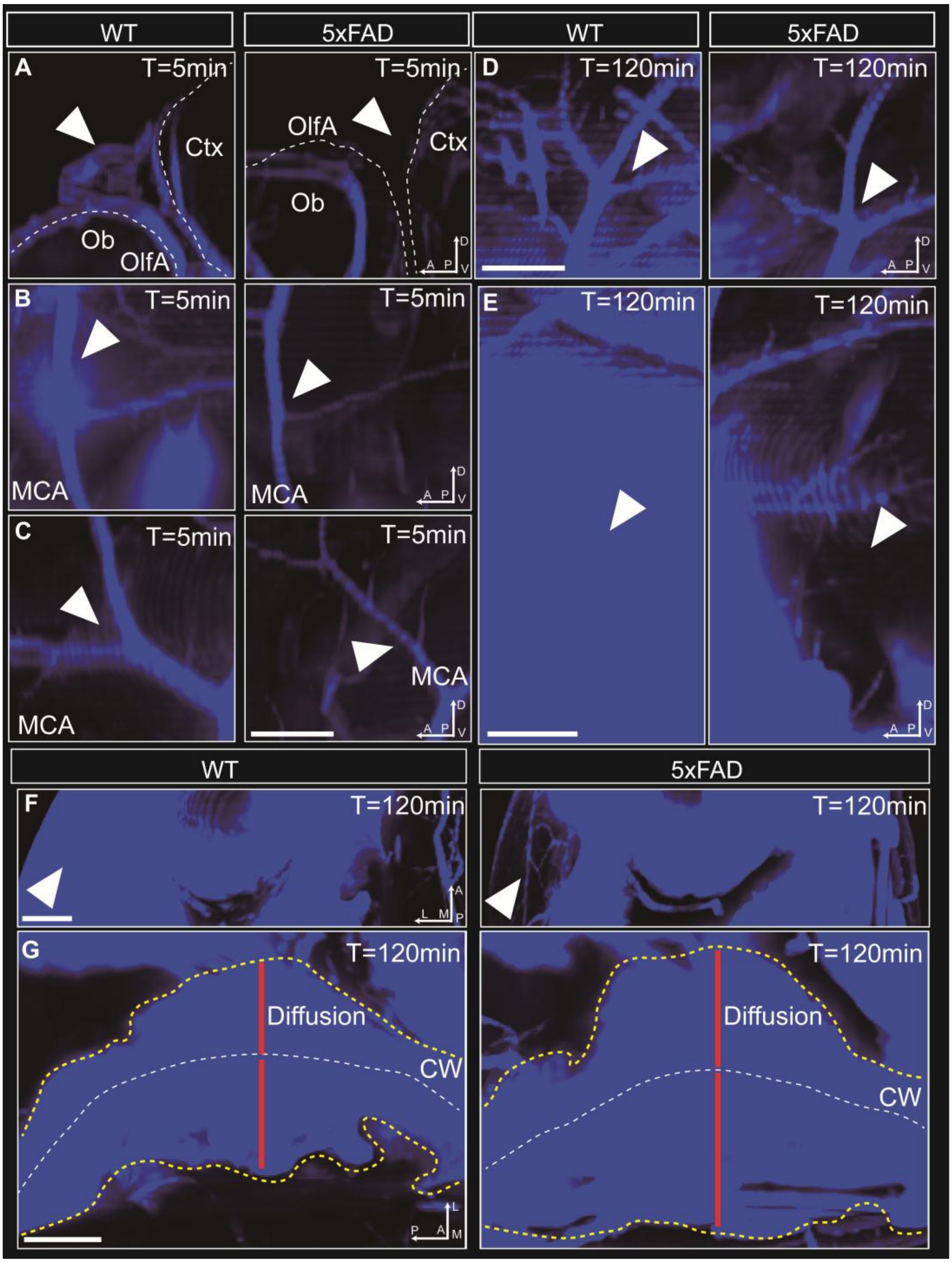


**Supplementary Figure. 2 | Three-dimensional light-sheet imaging reveals impaired dorsal tracer penetration and extensive ventral basal pooling in 5xFAD mice.** High-resolution 3D reconstructions of tissue-cleared hemibrains illustrating the volumetric distribution of the Evans Blue (EB) tracer at 5 min and 120 min post-intra-cisterna magna (I.C.M.) injection. Spatial orientations are indicated by axes: Anterior-Posterior (A-P), Dorsal-Ventral (D-V), and Medial-Lateral (M-L). **(A)** High-magnification views of the dorsal interface between the olfactory bulb (Ob) and cortex (Ctx) at 5 min post-injection. Regional boundaries are outlined by white dashed lines. Compared to wild-type (WT) controls, the 5xFAD group exhibits a visibly weaker EB signal (white arrowheads) propagating along the olfactory artery (OlfA) into this dorsal territory. **(B, C)** Magnified views of the middle cerebral artery (MCA) tract at 5 min, captured near its ventral origin (**B**) and more dorsally (**C**). White arrowheads highlight the MCA architecture. In both the proximal (ventral) and distal (dorsal) segments of the MCA, the 5xFAD mice display a markedly reduced EB signal intensity and diminished surrounding tissue diffusion compared to WT mice. **(D, E)** Extended 3D views of the dorsal MCA branches and the surrounding cortical parenchyma at 120 min. White arrowheads denote corresponding anatomical cortical regions in both groups. The WT brain exhibits robust and deep parenchymal EB penetration. In stark contrast, the 5xFAD cortex shows a near-complete absence of the EB tracer signal, confirming a severe failure in delayed dorsal fluid distribution. **(F)** Lateral profile views of the dorsal cortex at 120 min. White arrowheads emphasize the cortical superficial layers. The WT brain maintains widespread cortical tracer accumulation, whereas the 5xFAD group shows severely depleted EB signals across the equivalent lateral-dorsal regions. **(G)** High-magnification views of the ventral base surrounding the Circle of Willis (CW) at 120 min. The anatomical trajectory of the CW is indicated by white dashed lines. Yellow dashed lines demarcate the boundary of abnormal tracer diffusion into the surrounding ventral tissue, with the absolute diffusion distance highlighted by solid red lines. The 5xFAD group demonstrates a massive, pathological expansion of tracer diffusion around the CW compared to the restricted pattern in the WT group, visually corroborating the severe fluid stagnation and basal pooling secondary to dorsal influx failure.

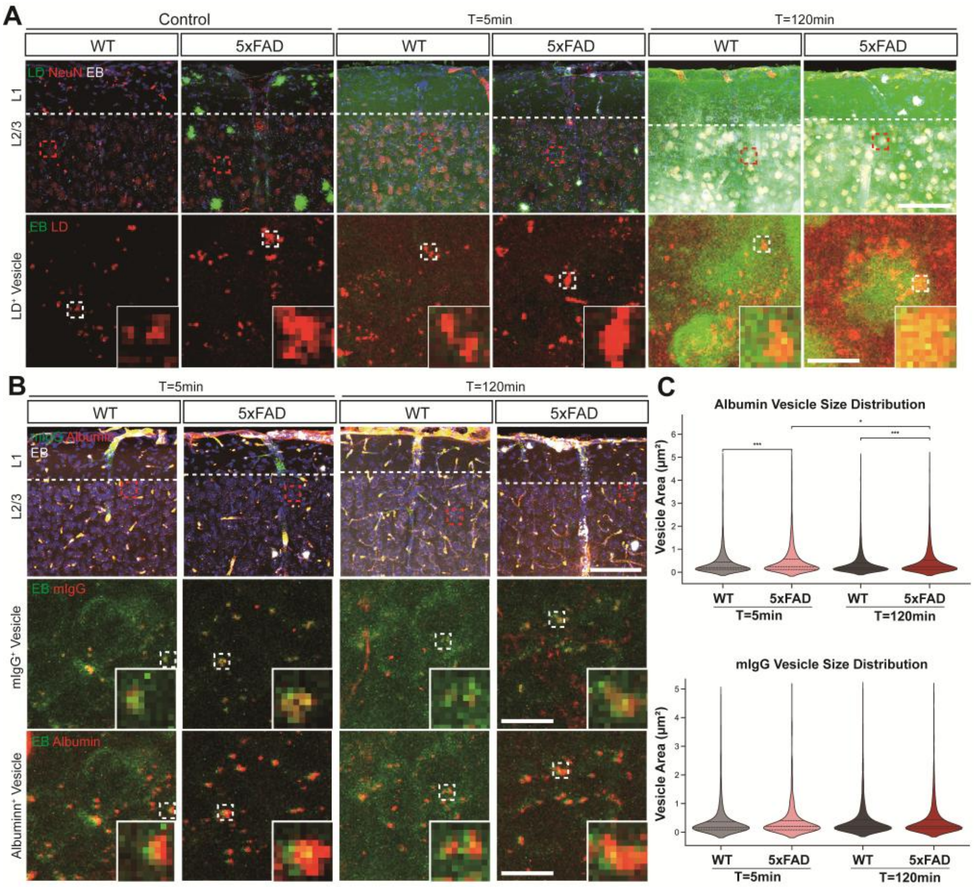


**Supplementary Figure. 3 | Internal cargo composition of vesicles and the effect of detergent-free tissue processing on tracer visualization. (A)** Representative confocal micrographs of the cortex (layers L1 and L2/3) in wild-type (WT) and 5xFAD mice at baseline (Control), 5 min, and 120 min following intra-cisterna magna (ICM) injection. Top: Merged images of Lipid droplet (LD, green), NeuN (red), and Evans Blue (EB, blue) (Scale bar, 100 μm). Bottom: Separated channel images of LD (red) and EB (green) at a 10x magnification (Scale bar, 10 μm). Insets display a 100x digital zoom of the dashed box regions. Note that the LD staining protocol intentionally omitted detergent-based permeabilization and washing steps. Consequently, parenchymal and soma-internalized EB signals appear as a diffuse blur rather than as sharply defined vesicular structures. Despite this methodological diffusion, the EB influx into the neuronal soma region remains distinctly higher than in the surrounding extracellular space. **(B)** Confocal imaging of the cortex in WT and 5xFAD mice at 5 min and 120 min post-ICM injection. Top: Merged images of mouse Immunoglobulin G (mIgG, green), Albumin (red), and EB (blue) (Scale bar, 100 μm). Middle and Bottom: Separated 10x magnified images displaying mIgG$^{+}$ vesicles (middle, red) and Albumin$^{+}$ vesicles (bottom, red) co-stained with EB (green) (Scale bars, 10 μm). Insets show a 100x magnification of the dashed box regions. In contrast to the peripheral exterior localization of CD31, AQP4, and ApoE, both mIgG and endogenous Albumin exhibit structural mixing and spatial overlap with the EB puncta. Furthermore, this internal co-localization of mIgG and Albumin with EB appears consistently maintained even after the vesicles are internalized within the neuronal soma. (C) Automated quantification of internal cargo vesicle size distributions. Violin plots illustrate the cross-sectional area ($\mu m^2$) of purely extravascular $Albumin^+$ (top) and $mIgG^+$ (bottom) vesicles in WT and 5xFAD mice at 5 min and 120 min post-injection. The internal dashed lines indicate the median and quartiles. Automated morphometric analysis revealed that 5xFAD mice exhibited a highly significant pathological increase in the size of $Albumin^+$ transport vesicles compared to WT controls at both 5 min (***$P < 0.001$) and 120 min (***$P < 0.001$) time points, suggesting an aberrant accumulation or swelling of the fluid-phase cargo core in the diseased state. In contrast, the size distribution of $mIgG^+$ vesicles remained relatively stable, showing no statistically significant variation between genotypes. Statistical significance was determined using one-way ANOVA followed by Tukey's post-hoc test (*$P < 0.05$, ***$P < 0.001$).

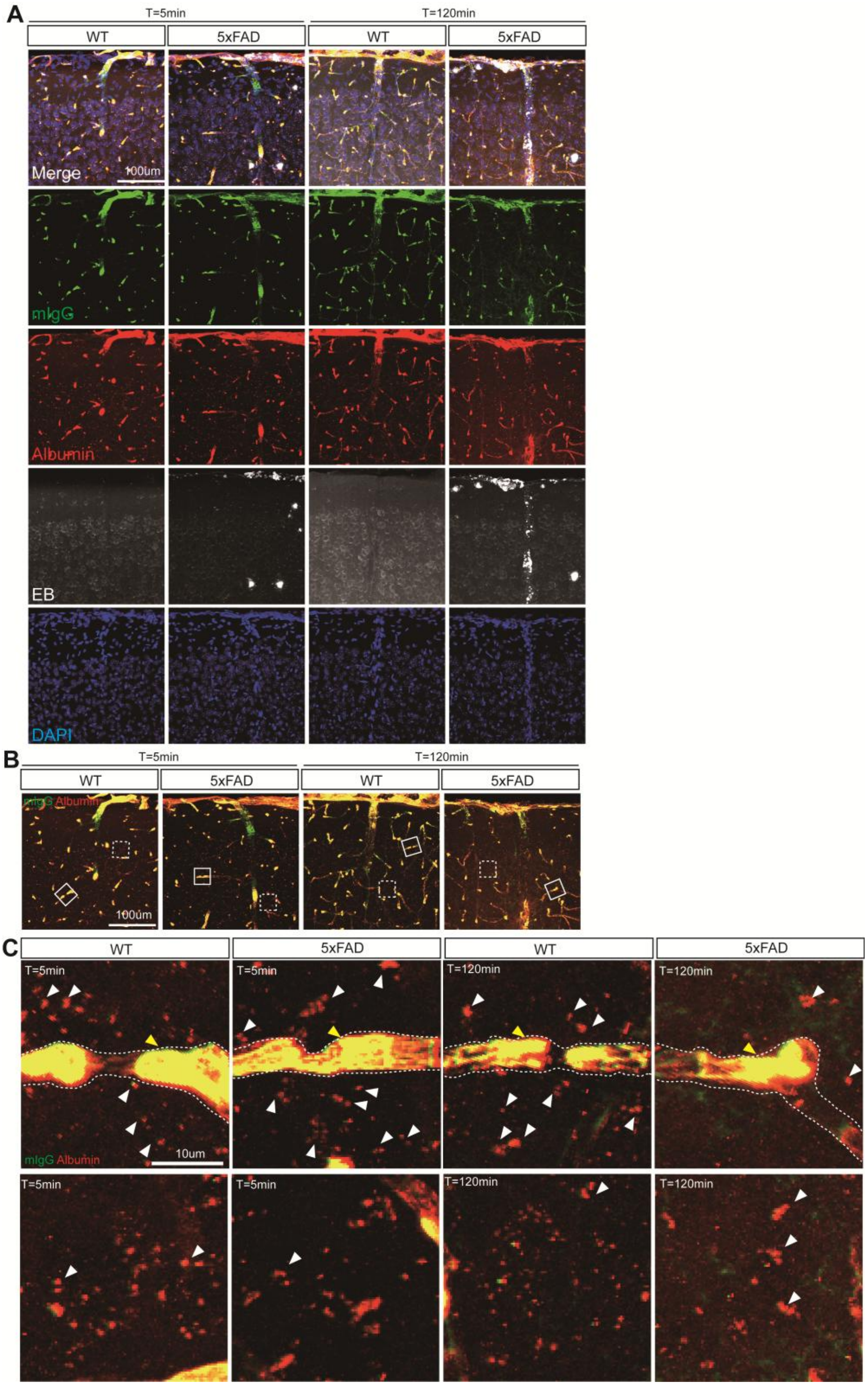


**Supplementary Figure. 4 | Endogenous albumin and mIgG colocalize within the cortical microvasculature and extravascular parenchymal vesicles. (A)** Representative high-resolution confocal micrographs of the cerebral cortex (layers L1 to L2/3) in wild-type (WT) and 5xFAD mice at 5 minutes and 120 minutes following intra-cisterna magna (I.C.M.) injection of the Evans Blue (EB) tracer. The top row displays merged composite images, while subsequent rows present isolated single-channel fluorescence signals: endogenous mouse immunoglobulin G (mIgG, green), endogenous Albumin (red), exogenous EB tracer (grayscale/white), and nuclei (DAPI, blue). **(B)** Dual-channel merged micrographs isolating mIgG (green) and Albumin (red) signals to evaluate the spatial relationship between these two major endogenous serum proteins within the neurovascular architecture. Solid white boxes indicate specific capillary regions targeted for high-magnification analysis. Scale bar, 100 μm. **(C)** High-magnification analyses of the capillary interfaces, derived from the solid white boxes in panel B. The capillary boundaries are delineated by white dashed lines. Across all groups (WT and 5xFAD at both 5 and 120 minutes post-injection), mIgG and albumin exhibit colocalization directly within the vascular lumen (yellow arrowheads). Furthermore, in the immediate extravascular space, discrete vesicular structures are observed traversing the parenchyma. These extravascular vesicles display colocalization of both mIgG and albumin (mIgG$^{+}$Albumin$^{+}$ puncta, indicated by white arrowheads). This suggests that endogenous macromolecular serum proteins may be co-packaged into shared transport vesicles operating within the brain parenchyma, a structural feature conserved across both physiological and pathological states. Scale bar, 10 μm.

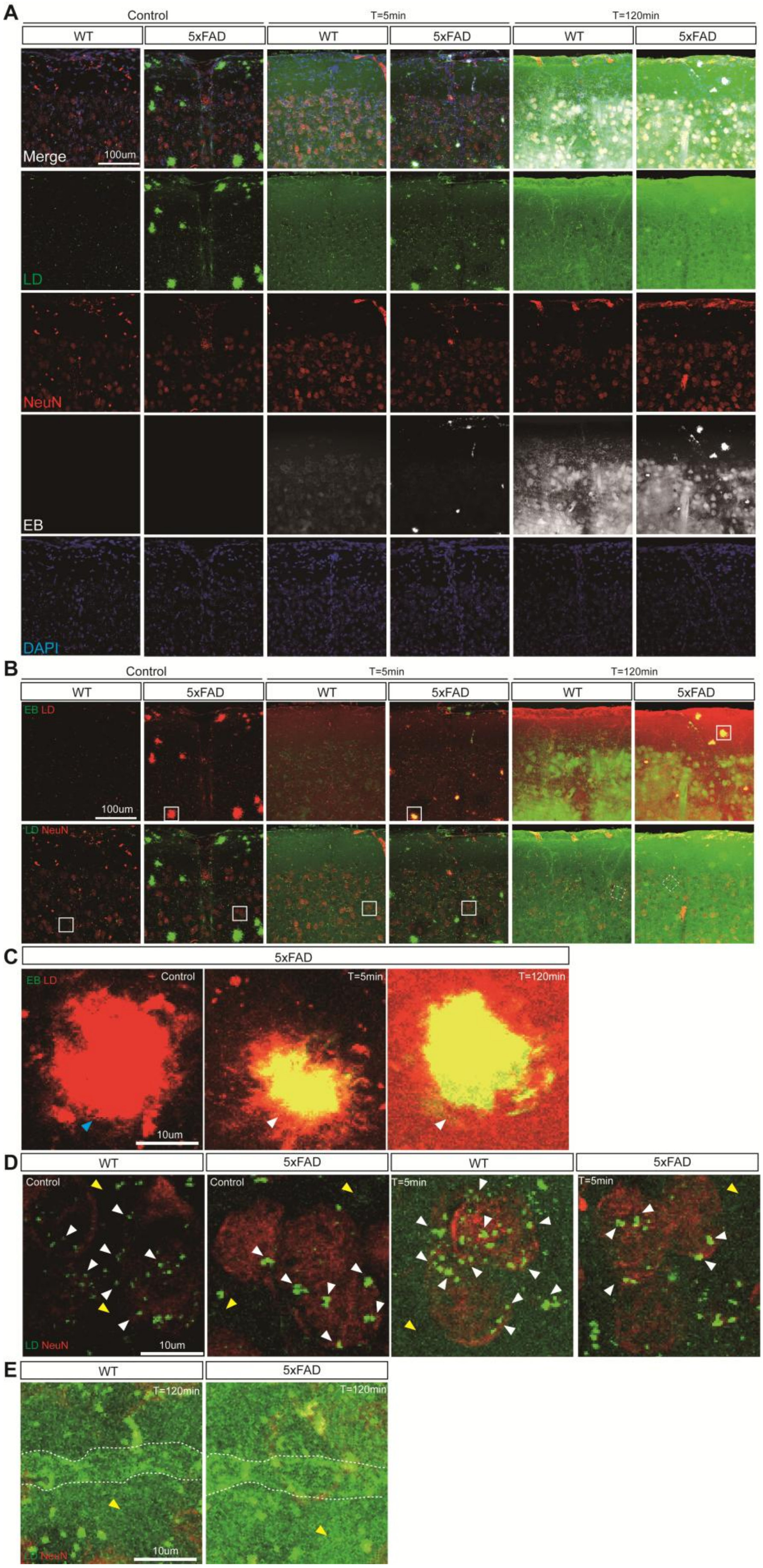
A
Control
T=5min
T=120min
WT
5xFAD
Merge
100um
LD
NeuN
EB
DAPI
B
EB LD
LD NeuN
C
5xFAD
10um
D
Control
T=5min
E
T=120min
10um

**Supplementary Figure. 5 | Spatiotemporal mapping of neuronal lipid droplet dynamics and pre-existing plaque lipidosis potentially driving CSF tracer entrapment.** **(A)** Representative high-resolution confocal micrographs of the cerebral cortex (layers L1 to L2/3) in wild-type (WT) and 5xFAD mice at baseline (Control), 5 minutes, and 120 minutes following intra-cisterna magna (I.C.M.) injection of the Evans Blue (EB) tracer. The top row displays merged composite images, while subsequent rows present isolated single-channel signals: Lipid Droplets (LD, green), neuronal soma (NeuN, red), EB tracer (grayscale/white), and nuclei (DAPI, blue). **(B)** Dual-channel merged micrographs isolating specific functional interactions. The top row (EB pseudo-colored in green; LD pseudo-colored in red) highlights the spatial relationship between the exogenous CSF tracer and the tissue lipid architecture. The bottom row (LD in green; NeuN in red) visualizes intracellular neuronal lipid dynamics and extracellular matrix (ECM) lipid distribution. Scale bars, 100 μm. **(C)** High-magnification zoomed views of amyloid plaques in 5xFAD mice (from the solid white boxes in the top row of panel b) resolving the interaction between LDs and the EB tracer. Notably, even prior to tracer injection (Control), massive LD aggregation is natively established at the amyloid plaques (blue arrowhead). Following I.C.M. injection, the EB tracer robustly co-aggregates with these pre-existing lipid-rich plaques at both 5 and 120 minutes (white arrowheads). This suggests that amyloid plaques are pathologically pre-loaded with lipids, potentially functioning as physical sinks that trap incoming CSF. Scale bar, 10 μm. **(D)** High-magnification views of layer L2/3 pyramidal neurons (from the solid white boxes in the bottom row of panel b) evaluating physiological vs. pathological neuronal lipid vesicle responses. At baseline (Control), discrete LD vesicles are present within the $NeuN^+$ neuronal soma of both WT and 5xFAD mice (white arrowheads), though 5xFAD mice exhibit elevated basal LD signals in the ECM (yellow arrowheads). Following CSF influx at 5 minutes, WT neurons mount a robust physiological response, characterized by an increase in both the size and number of intracellular lipid vesicles. In contrast, 5xFAD neurons exhibit a blunted lipid response, failing to significantly alter their internal vesicular pool compared to baseline. Furthermore, the dynamic increase of LDs in the ECM observed in WT mice appears abnormally restricted in the 5xFAD group during this early phase. Scale bar, 10 μm. **(E)** High-magnification analysis of the vascular interface and ECM at 120 minutes post-injection. By this late clearance phase, both WT and 5xFAD groups exhibit pronounced LD accumulation along the capillary boundaries (delineated by white dashed lines). However, the 5xFAD cortex demonstrates a pathological exacerbation of LD signaling pooling within the ECM space (yellow arrowhead), reflecting an apparent macroscopic failure to clear fluid-mobilized lipids from the parenchymal environment. Scale bar, 10 μm.

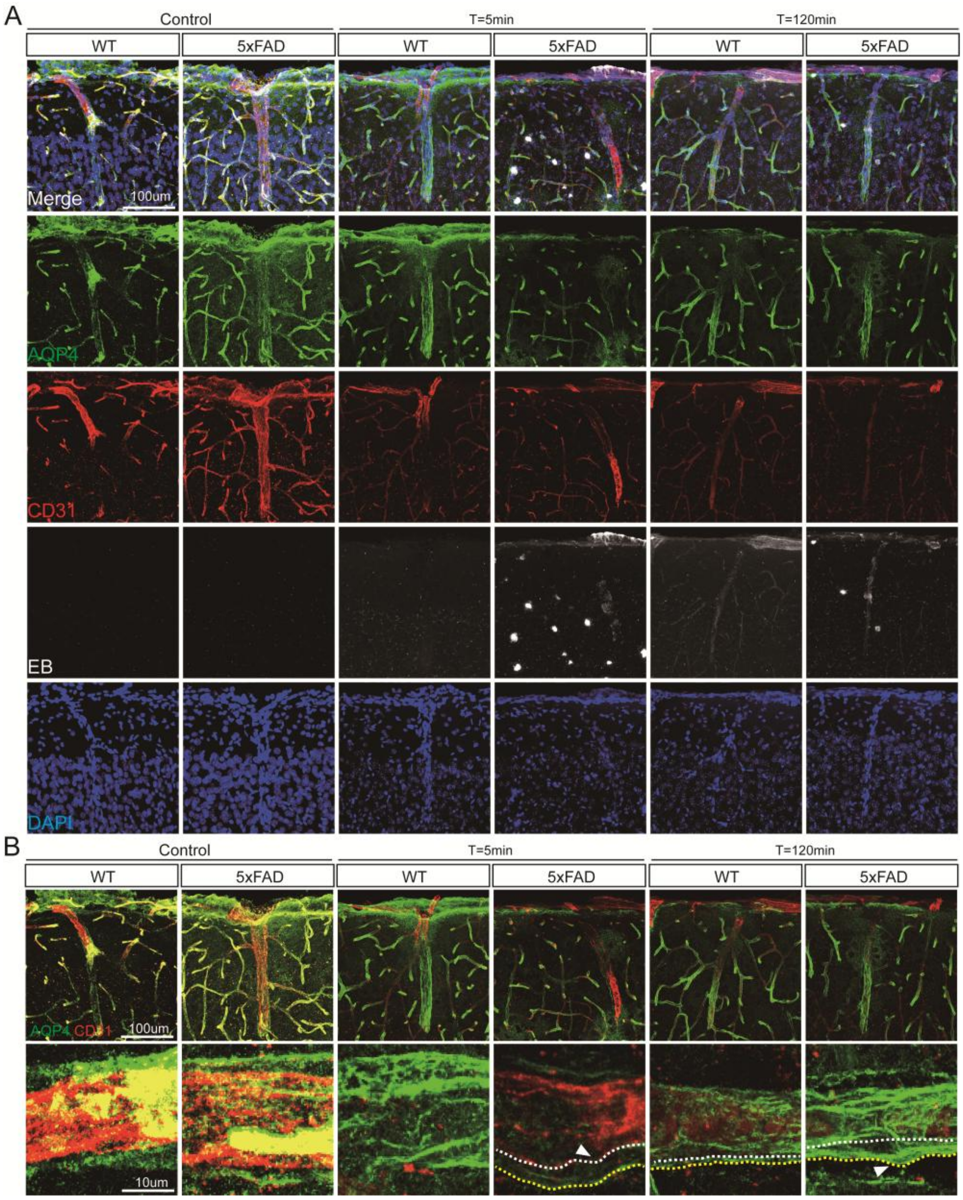


**Supplementary Figure. 6 | Single-channel resolution of the gliovascular interface and structural expansion of the perivascular boundary in the Alzheimer's disease cortex. (A)** Representative high-resolution confocal micrographs of the cerebral cortex (layers L1 to L2/3) in wild-type (WT) and 5xFAD mice at baseline (Control), 5 minutes, and 120 minutes following intra-cisterna magna (I.C.M.) injection of the Evans Blue (EB) cerebrospinal fluid tracer. The top row displays the merged composite images illustrating the spatial relationship between the vasculature, astrocytic boundary, and tracer dynamics. The subsequent rows present the isolated, unmerged single-channel fluorescence signals: aquaporin-4 (AQP4, green) delineating the astroglial endfoot boundary; CD31 (red) marking the vascular endothelium; EB (grayscale/white) tracking the distribution of the CSF tracer; and DAPI (blue) identifying cell nuclei. Visualization of the separated channels confirms the gross anatomical architecture. The isolated EB channel reveals fluid dynamics in 5xFAD mice: tracer accumulation strictly confined within the AQP4 boundary (perivascular stagnation) and focal tracer aggregations in the deep parenchyma (plaque-associated trapping), which become exacerbated over time. **(B)** Dual-channel visualization and high-magnification analysis highlighting the structural characteristics of the gliovascular interface. The top row displays the merged AQP4 (green) and CD31 (red) signals across matching time points (Scale bar, 100 μm). The bottom row provides zoomed-in views of the cortical penetrating vessels (Scale bar, 10 μm). To assess the perivascular space (PAS), the vascular endothelial boundary is delineated by white dashed lines, and the astroglial endfoot boundary is delineated by yellow dashed lines. In WT mice, the AQP4 and CD31 layers remain tightly apposed, maintaining a narrow PAS. Conversely, in 5xFAD mice, structural decoupling is observed. White arrowheads highlight the pathological widening of the distance between the CD31 and AQP4 boundaries at 5 and 120 minutes post-injection. This expansion of the PAS correlates with the perivascular stagnation of the EB tracer seen in panel a, suggesting a structural breakdown of the gliovascular architecture in the AD model.

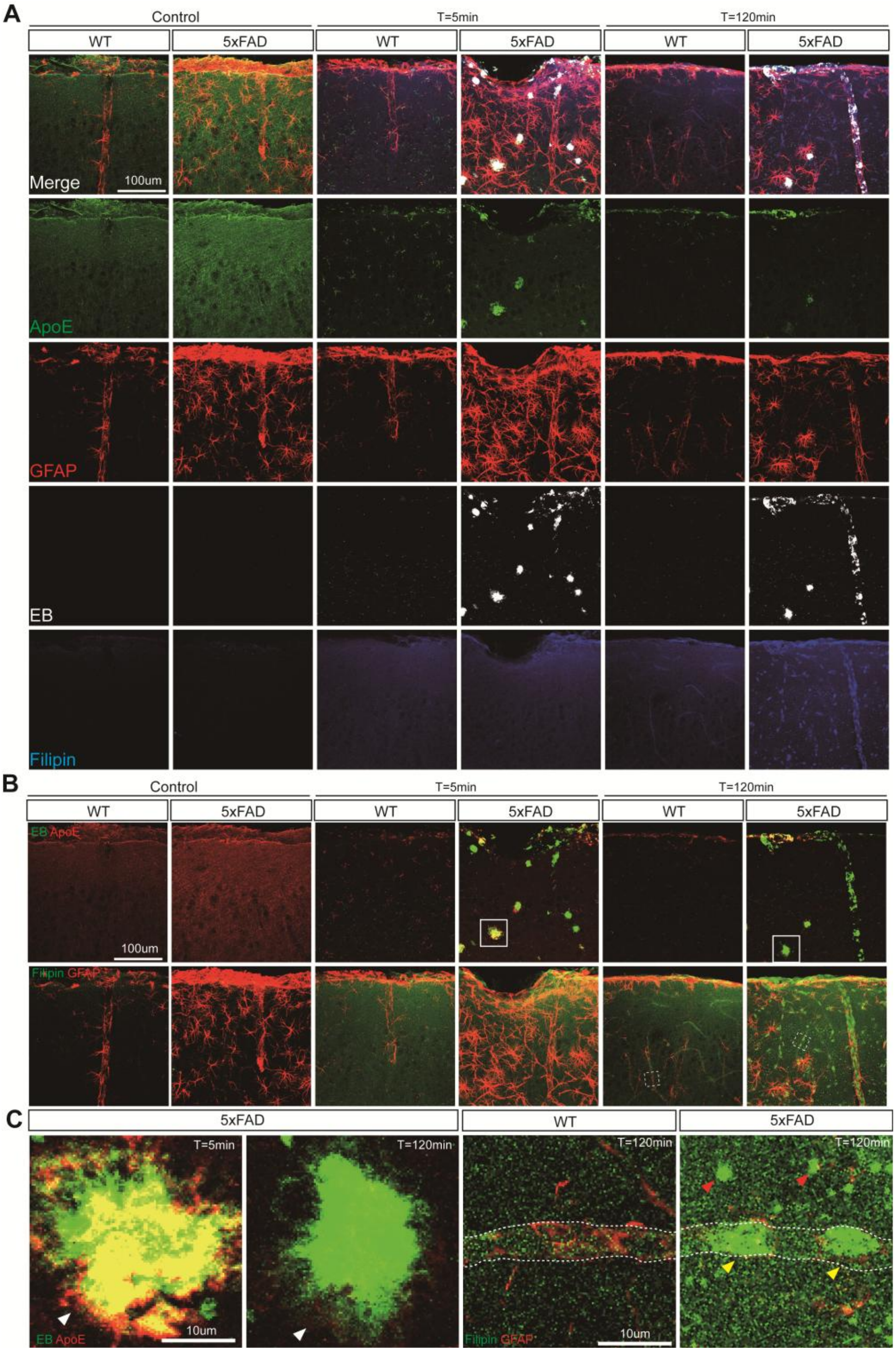


**Supplementary Figure. 7 | Spatiotemporal dynamics of ApoE-mediated lipid transport, plaque-associated CSF trapping, and perivascular cholesterol retention. (A)** Representative high-resolution confocal micrographs of the cerebral cortex (layers L1 to L2/3) in wild-type (WT) and 5xFAD mice at baseline (Control), 5 minutes, and 120 minutes following intra-cisterna magna (I.C.M.) injection of the Evans Blue (EB) cerebrospinal fluid tracer. The top row displays merged images, while subsequent rows present isolated single-channel signals: ApoE (green), GFAP (red), EB (grayscale/white), and

Filipin (blue). At baseline, isolated GFAP and ApoE channels demonstrate astrogliosis and focal ApoE deposition in 5xFAD mice. Post-injection, the isolated EB channel tracks the temporal progression of parenchymal fluid distribution in WT mice, contrasting with the focal tracer trapping observed in the 5xFAD brain. Scale bar, 100 μm. **(B)** Dual-channel merged micrographs isolating specific molecular intersections to evaluate functional lipid and tracer dynamics. The top row (EB in red, ApoE in green) highlights the interaction between the exogenous CSF tracer and endogenous lipid transport proteins. The bottom row (Filipin in green, GFAP in red) visualizes the spatial relationship between cholesterol content and the reactive astrocytic network. Scale bar, 100 μm. **(C)** High-magnification analyses of specific microenvironments, resolving cellular patterns of lipid and fluid clearance. Left: Zoomed-in views of the solid white boxes in panel b, capturing EB/ApoE dynamics. Prior to injection, ApoE appears diffusely distributed in the parenchyma. At 5 minutes post-injection, ApoE exhibits cellular internalization into astrocytes in WT mice. In contrast, 5xFAD mice exhibit co-aggregation of ApoE and the EB tracer directly at amyloid plaques (white arrowheads), a fluid-trapping event that persists through 120 minutes. Right: Zoomed-in views of the dashed white boxes in panel B, focusing on cholesterol (Filipin) dynamics at the capillary interface at 120 minutes post-injection. While overall Filipin signals elevate post-injection, their clearance trajectories diverge. In WT mice, Filipin accumulates within the vascular lumen, indicating perivascular lipid efflux. However, in 5xFAD mice, luminal Filipin appears reduced, and the cholesterol signal is retained within the parenchyma. Focal Filipin aggregation occurs within perivascular cells morphologically consistent with pericytes (yellow arrowheads) tightly enveloping the capillaries (outlined by white dashed lines), suggesting alterations in perivascular cholesterol clearance and cellular lipid stress in the AD neurovascular unit. Scale bar, 10 μm.

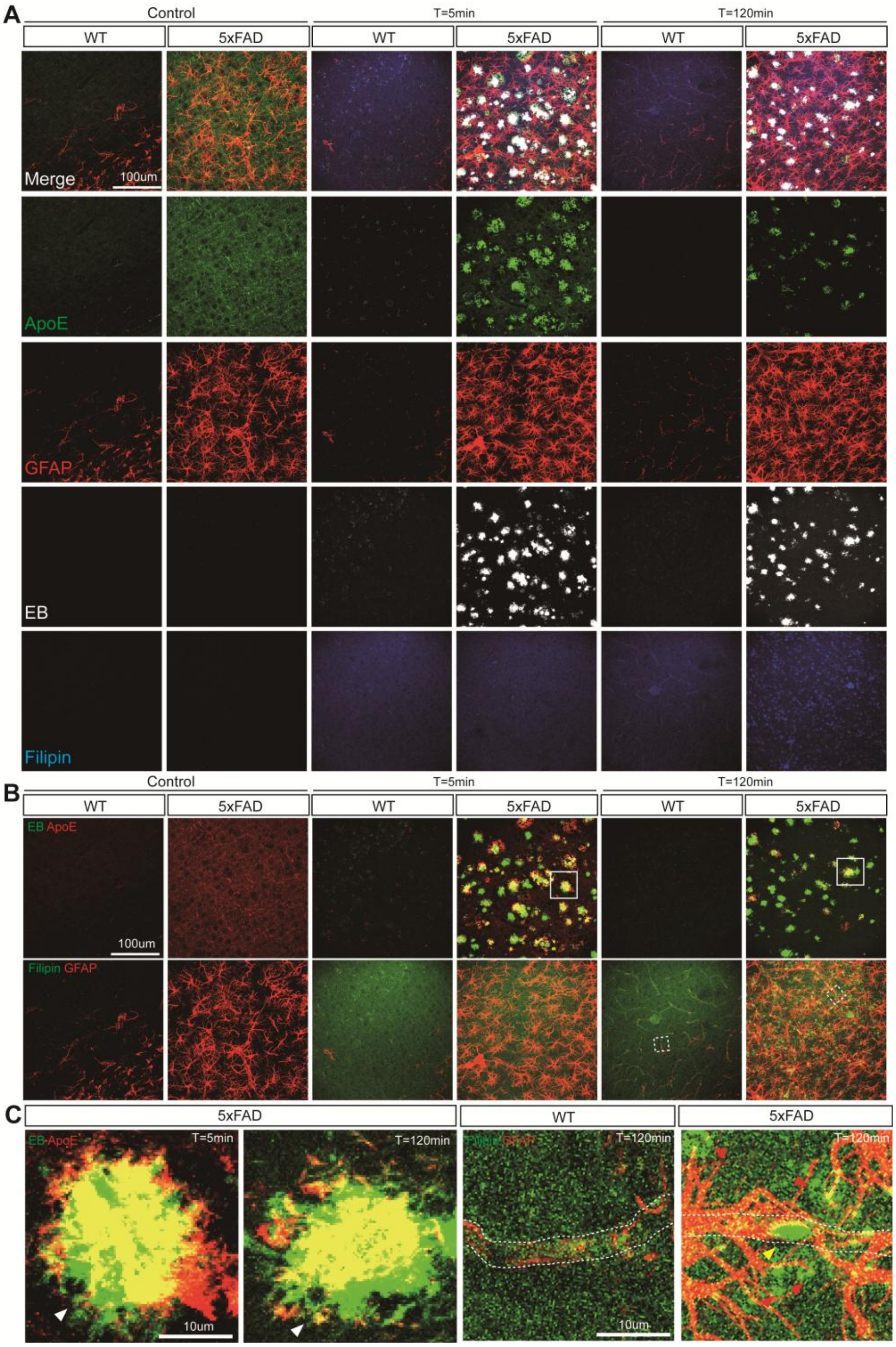


**Supplementary Figure. 8 | Deep cortical (Layer 6) dynamics reveal fluid-induced ApoE/tracer trapping at amyloid plaques and pericytic cholesterol retention. (A)** Representative high-resolution confocal micrographs of the deep cerebral cortex (Layer 6) in wild-type (WT) and 5xFAD mice at baseline (Control), 5 minutes, and 120 minutes following intra-cisterna magna (I.C.M.) injection of the Evans Blue (EB) cerebrospinal fluid tracer. The top row displays merged images, while subsequent rows show isolated single-channel signals: ApoE (green), GFAP (red), EB (grayscale/white), and Filipin (blue). Layer 6 is a region characterized by a high density of amyloid plaques in the AD model. At baseline, isolated ApoE remains diffusely distributed without forming focal aggregations at plaques in 5xFAD mice, while astrogliosis (GFAP) is prominent. Following injection, the WT group demonstrates a diffuse tracer distribution. In contrast, the influx of the CSF tracer in 5xFAD mice is associated with localized EB trapping. Scale bar, 100 μm. **(B)** Dual-channel merged micrographs isolating specific functional intersections. The top row (EB in red, ApoE in green) captures the interaction between the exogenous CSF tracer and endogenous ApoE. The bottom row (Filipin in green, GFAP in red) visualizes cholesterol clearance trajectories in the context of the reactive astrocytic network. Scale bar, 100 μm. **(C)** High-magnification analyses resolving deep-layer cellular patterns of fluid and lipid distribution. Left: Zoomed-in views of the solid white boxes in panel B, detailing EB/ApoE dynamics at amyloid plaques. While ApoE does not spontaneously aggregate at plaques prior to injection, the physical arrival of the CSF tracer appears to act as a catalyst. At 5 and 120 minutes post-injection, the 5xFAD model exhibits fluid-induced co-aggregation of ApoE and EB directly at the amyloid plaques (white arrowheads). This suggests that plaques function as physical sinks, capturing lipid-transport proteins upon bulk fluid flow. Right: Zoomed-in views of the dashed white boxes in panel b, highlighting cholesterol (Filipin) dynamics at the capillary interface at 120 minutes. While both groups show elevated Filipin signals post-injection, their spatial distribution diverges. In WT mice, Filipin accumulates within the vascular lumen, indicating perivascular lipid efflux. Conversely, in 5xFAD mice, luminal Filipin appears largely absent, and the cholesterol signal is retained within the parenchyma amidst reactive astrocytes. Consistent with the pathology observed in superficial cortical layers, focal Filipin aggregation occurs within perivascular cells anatomically corresponding to pericytes (yellow arrowheads) ensheathing the capillaries (outlined by white dashed lines), indicating that alterations in perivascular cholesterol clearance and pericytic lipid stress are conserved across the cortical depth in the AD brain. Scale bar, 10 μm.

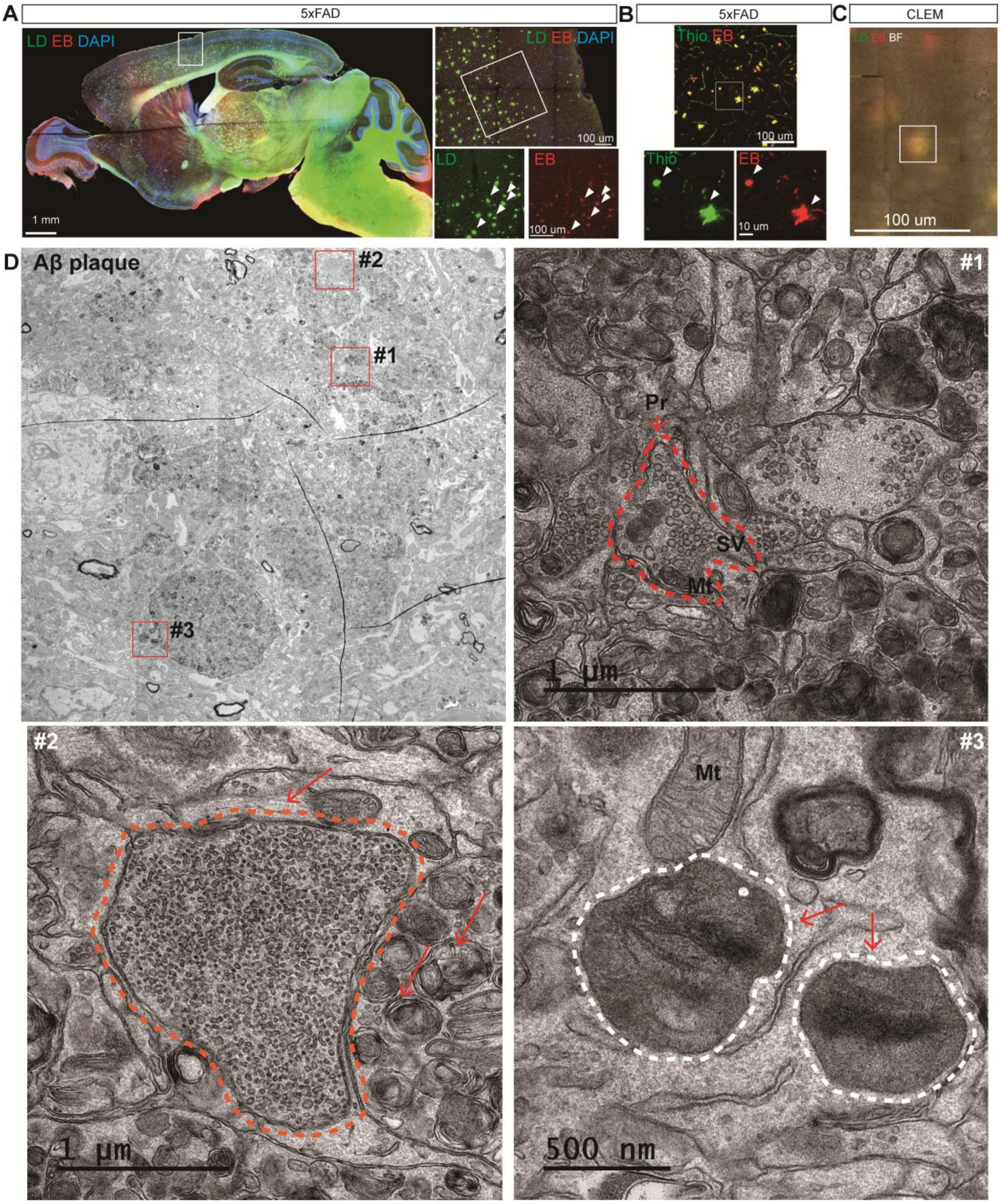


**Supplementary Figure. 9 | Ultrastructural characterization suggests CSF stagnation and lipid co-aggregation at the amyloid plaque microenvironment in the AD brain. (A)** Representative whole-brain sagittal section imaging of a 5xFAD mouse post-intra-cisterna magna (ICM) injection of Evans Blue (EB) tracer. Left: Merged panoramic view of Lipid droplet (LD, green), EB (red), and DAPI (blue) (Scale bar, 1 mm). Top right: Magnified view of the solid white box region from the left panel, showing the cortical distribution (Scale bar, 100 μm). Bottom right: Separated single-channel images of the solid white boxed area in the top right panel. White arrowheads highlight the spatial colocalization of LD and

EB signals, suggesting that the stagnated CSF may be enriched with lipids (Scale bar, 100 μm). **(B)** Confocal imaging evaluating the distribution of CSF at amyloid plaques in the 5xFAD cortex. Top: Merged image of amyloid plaques stained with Thioflavin S (Thio, green) and EB (red) (Scale bar, 100 μm). Bottom: Separated, high-magnification channel images of the solid white box region. White arrowheads indicate the structural colocalization of Thio-positive plaques and EB tracer, indicating that amyloid plaques appear to function as major aggregation sites for disrupted CSF flow (Scale bar, 10 μm). **(C)** Correlative light and electron microscopy (CLEM) linking fluorescence signals to nanoscale architecture. The confocal fluorescence image (LD, green; EB, red; overlaid with brightfield, BF) is aligned with the corresponding transmission electron microscopy (TEM) micrograph to target the specific amyloid plaque structure associated with capturing the EB/LD aggregates (Scale bar, 100 μm). **(D)** Representative TEM micrograph of the targeted amyloid plaque microenvironment identified in C. The large overview image displays the complex pathological architecture of the plaque area. Dashed red boxes (#1–3) indicate regions of interest that are magnified in the adjacent panels. The high-magnification panels reveal ultrastructural features, including dystrophic neurites, synaptic vesicles (SV), mitochondria (Mt), and electron-dense vesicular and lipid structures (red dashed outlines and arrows) intimately associated with the plaque core (Scale bars, 1 μm and 500 nm, as indicated).

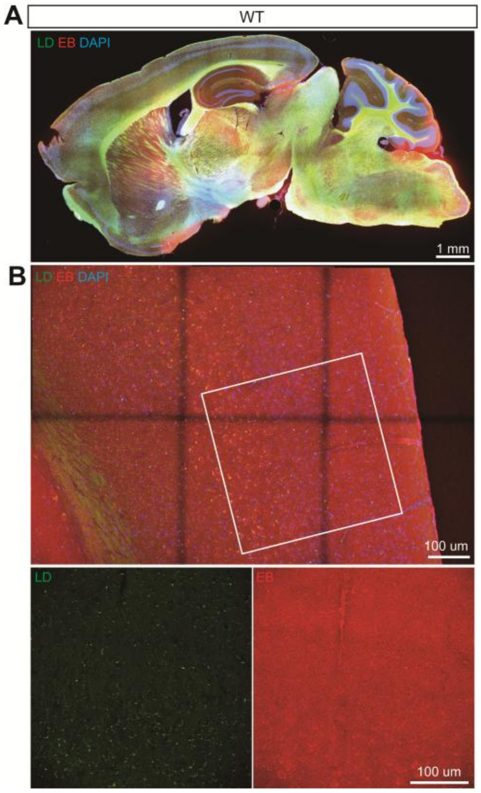


**Supplementary Figure. 10 | Macroscopic and regional mapping of lipid droplets and CSF tracer distribution in the wild-type brain. (A)** Representative panoramic imaging of a whole-brain sagittal section from a wild-type (WT) mouse acquired via inverted confocal microscopy. The tissue was harvested following intra-cisterna magna (I.C.M.) injection of the Evans Blue (EB) tracer. The merged image displays the baseline distribution of Lipid Droplets (LD, green), the EB cerebrospinal fluid tracer (red), and cell nuclei (DAPI, blue). Scale bar, 1 mm. **(B)** High-resolution mesoscopic evaluation of the cerebral cortex to establish the structural baseline for subsequent correlative light and electron microscopy (CLEM). The top panel displays a merged, tiled overview of the cortical architecture. The solid white box demarcates the targeted superficial cortex (layers L1 to L3). The bottom panels present the isolated single-channel fluorescence signals from this specific region: LD (left, green) highlighting discrete lipid vesicles within the perivascular space and neuronal soma, and EB (right, red) illustrating the diffuse permeation of the tracer throughout the healthy parenchyma. Scale bars, 100 µm.

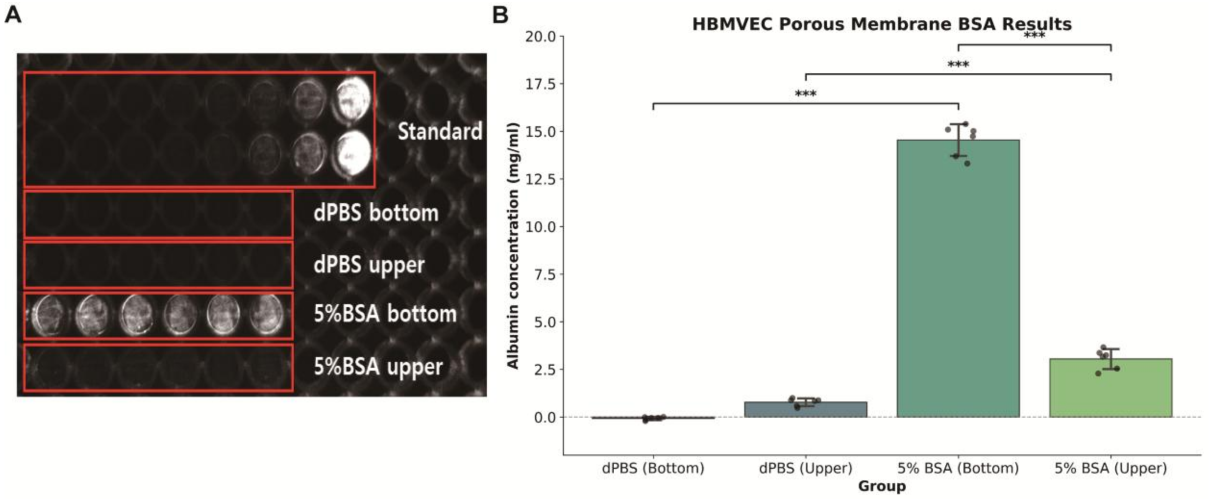


**Supplementary Figure. 11 | Human brain microvascular endothelial cells (HBMVECs) actively transcytose albumin, providing a potential molecular substrate for CEPT vesicle formation. (A)** Representative macroscopic fluorescence image of the in vitro microplate assay designed to evaluate endothelial protein permeability. HBMVECs were cultured on porous Transwell membranes to model the blood-brain barrier interface. The apical (upper) chamber was treated with either dPBS (vehicle control) or 5% bovine serum albumin (BSA). Following the transcytosis incubation period, media from both the upper and basolateral (bottom) chambers were collected, incubated with the Evans Blue (EB) tracer, and imaged. The top two rows display the serially diluted albumin standard curve. The subsequent rows display the experimental groups: dPBS bottom, dPBS upper, 5% BSA bottom, and 5% BSA upper. Consistent with the tracer's binding properties, fluorescence emission is visible in the 5% BSA-treated groups, confirming the presence of albumin. **(B)** Quantitative microplate reader analysis of albumin concentration (mg/mL) across the experimental groups, interpolated directly from the standard curve. In the dPBS control groups, albumin concentrations were near zero in both chambers. Following 5% BSA administration to the apical chamber, an accumulation of albumin was detected in the basolateral compartment (5% BSA Bottom; ~14.5 mg/mL), exceeding the residual concentration in the apical chamber (5% BSA Upper; ~3.0 mg/mL) (***$P < 0.001$). This quantitative result demonstrates that HBMVECs can transcytose bulk albumin across the endothelial barrier layer.

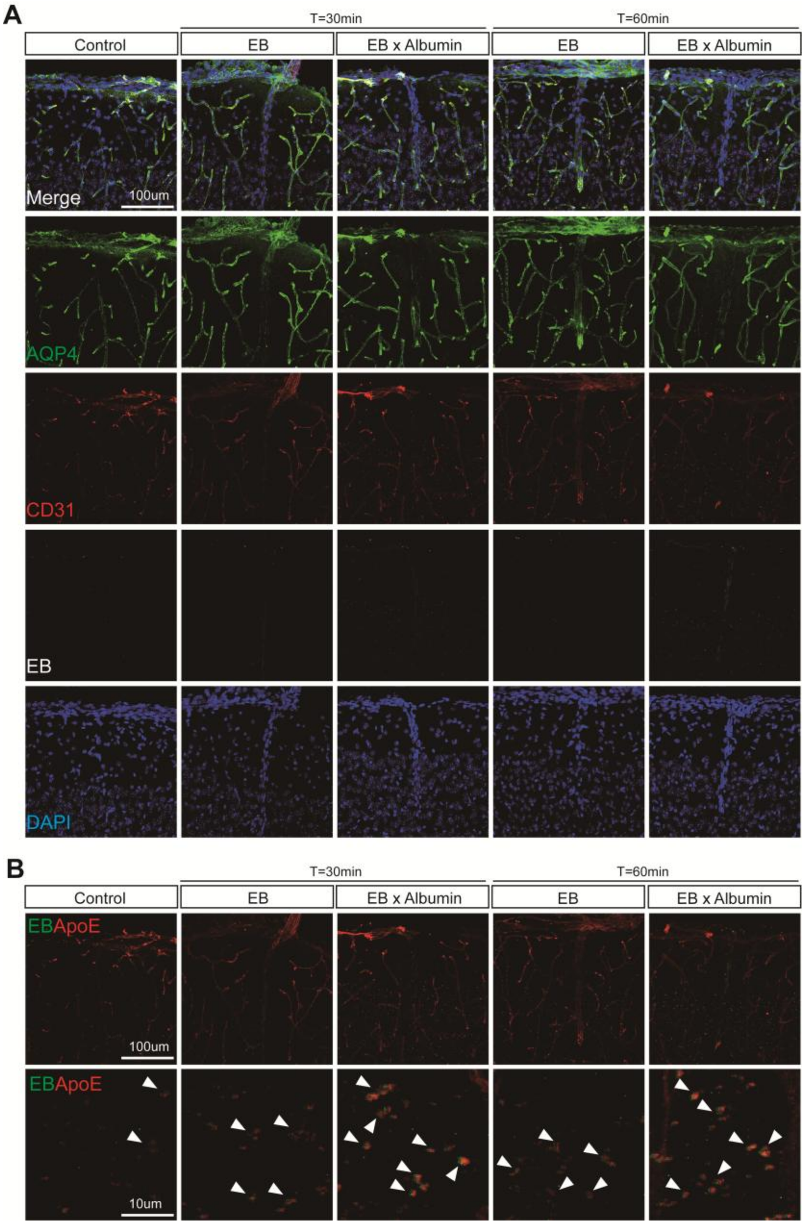


**Supplementary Figure. 12 | Endothelial-transcytosed albumin is associated with enhanced formation and parenchymal infiltration of $CD31^{+}$ vesicles following systemic administration.** **(A)** Representative high-resolution confocal micrographs of the cerebral cortex in wild-type (WT) mice at baseline (Control) and at 30 and 60 minutes following intra-retro-orbital (I.R.O.) injection. To evaluate potential molecular triggers of systemic tracer uptake, mice were injected with either Evans Blue alone (EB) or an EB-Albumin premix (EB x Albumin). To mimic physiological blood-brain barrier processing, the EB x Albumin complex was prepared by pre-conditioning albumin through a human brain microvascular endothelial cell (HBMVEC) porous membrane transcytosis assay (albumin administered to the apical chamber, with basolateral media collected and premixed with EB) prior to I.R.O. injection. The top row displays merged composite images, followed by isolated single-channel signals: AQP4 (green), CD31 (red), EB (grayscale/white), and DAPI (blue). Scale bar, 100 µm. **(B)**

Dual-channel merged micrographs isolating CD31 (red) and EB (green) signals to visualize systemic uptake into parenchymal vesicles. The top row displays overview images (Scale bar, 100 μm), and the bottom row provides high-magnification views (10x zoom) of the parenchymal space (Scale bar, 10 μm). In the baseline Control group, native CD31 vesicular structures (white arrowheads) are observed distributed within the brain parenchyma prior to injection. Following I.R.O. injection of EB, the abundance of these vesicles appears increased, with the systemic EB tracer co-localizing within them ($CD31^{+}EB^{+}$). Systemic administration of the HBMVEC-transcytosed EB x Albumin premix corresponds to an increase in the size and quantity of these $CD31^{+}EB^{+}$ vesicles compared to EB alone. This enhanced vesicular formation and tracer incorporation persist from 30 to 60 minutes post-injection, suggesting that endothelial-processed albumin may act as a molecular factor for facilitating CEPT vesicular transport from the systemic circulation into the deep brain parenchyma.

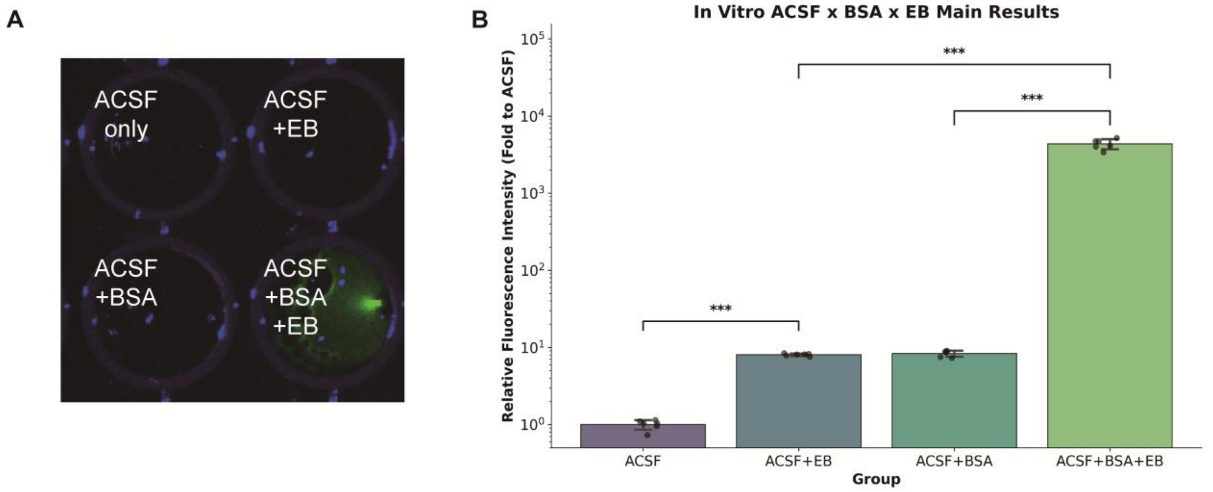


**Supplementary Figure. 13 | Robust fluorescence emission of Evans Blue appears dependent on albumin binding. (A)** Representative macroscopic fluorescence imaging (ChemiDoc) of in vitro microplate wells containing artificial cerebrospinal fluid (ACSF) alone, ACSF with Evans Blue (ACSF+EB), ACSF with bovine serum albumin (ACSF+BSA), and ACSF with both BSA and EB (ACSF+BSA+EB). In the absence of albumin, the EB tracer (ACSF+EB) exhibits minimal fluorescence emission, appearing comparable to blank controls. Conversely, the physical binding of EB to albumin (ACSF+BSA+EB) triggers a distinct fluorescence signal. **(B)** Quantitative microplate reader analysis of relative fluorescence intensity (fold change to ACSF only) across the four experimental groups. The quantitative data corroborate the visual imaging: while ACSF+EB and ACSF+BSA yield baseline signals (approximately 8.06-fold and 8.35-fold relative to ACSF alone, respectively), the formation of the EB-albumin complex (ACSF+BSA+EB) results in an amplification of fluorescence intensity (>4,365-fold relative to ACSF; ***$P < 0.001$).
